# Updated taxonomy of the family *Rhizobiaceae* with proposals for 10 novel genera and 35 novel combinations

**DOI:** 10.1101/2025.11.20.689363

**Authors:** Natalia Naranjo-Robayo, Tia L. Harrison, Oona Esme, Esther Menéndez, Nemanja Kuzmanović, Álvaro Peix, J. Peter W. Young, George C. diCenzo

## Abstract

The family *Rhizobiaceae* of the class *Alphaproteobacteria* is highly diverse and currently consists of at least 276 validly published or proposed species across 38 genera. Despite several recent studies proposing revisions to the family *Rhizobiaceae*, anomalies and inconsistencies in the taxonomy of this family remain. Here, we revisit the taxonomy of the family *Rhizobiaceae* with a focus on the genus *Rhizobium*. First, we generated whole genome sequences for 12 *Rhizobium* type strains that previously lacked publicly available genome sequences. We then applied an established phylogenomic framework to reappraise the taxonomic classification of 242 *Rhizobiaceae* type strains. Our data suggest that *Rhizobium aegyptiacum* is a later heterotypic synonym of *Rhizobium aethiopicum*, and they contradict a recent suggestion that *Rhizobium azibense* and *Rhizobium gallicum* are synonymous. In addition, we propose the formation of ten new genera (*Allohoeflea* gen. nov., *Arminia* gen. nov., *Fluviimicrobium* gen. nov., *Gillisella* gen. nov., *Limnomicrobium* gen. nov., *Martinezia* gen. nov., *Neohoeflea* gen. nov., *Parahoeflea* gen. nov., *Velazquezia* gen. nov., and *Yannia* gen. nov.) and 35 novel combinations to fix paraphyletic genera or account for monophyletic type strains that are clearly distinguishable based on core-proteome average amino acid identity (cpAAI) comparisons. Lastly, our data suggest that the type strain of *Rhizobium arsenicireducens* may have been lost, and that either a neotype should be designated or the taxonomic status of this species should be revised.

## INTRODUCTION

The family *Rhizobiaceae* of the class *Alphaproteobacteria* was proposed in 1938 and includes many rhizobia (nitrogen-fixing legume symbionts), agrobacteria (which induce tumours on diverse plant species), and other environmental bacteria [1, 2]. Currently, the family *Rhizobiaceae* consists of at least 276 validly published or proposed species across the following 38 genera: *Affinirhizobium*, *Agrobacterium*, *Aliirhizobium*, “*Allopararhizobium*”, *Allorhizobium*, *Ciceribacter*, “*Ectorhizobium*”, *Endobacterium*, *Ensifer*, *Ferirhizobium*, *Ferranicluibacter*, *Flavimaribacter*, *Gellertiella*, *Georhizobium*, *Heterorhizobium*, *Hoeflea*, *Lentilitoribacter*, *Liberibacter*, *Martelella*, “*Metarhizobium*”, *Mycoplana*, *Neopararhizobium*, *Neorhizobium*, *Onobrychidicola*, “*Oryzifoliimicrobium*”, *Paenirhizobium*, “*Paramartelella*”, *Pararhizobium*, *Peteryoungia*, “*Candidatus* Porivivens”, *Pseudohoeflea*, *Pseudorhizobium*, “*Candidatus* Reichenowia”, *Rhizobium*, *Shinella*, *Sinorhizobium*, *Terrirhizobium*, and *Xaviernesmea* [3, 4].

Recent years have seen multiple studies proposing revisions to the family *Rhizobiaceae* to fix known or newly revealed taxonomic inconsistencies [5–7]; however, anomalies in the taxonomy of this family remain. One challenge in revising the taxonomy of the family *Rhizobiaceae* is that many species were described before whole genome sequencing was as accessible as it is nowadays and, as a result, many type strains continue to lack publicly available genome sequences. Due to this, the taxonomic classification of many species currently cannot be addressed using state-of-the art phylogenomic approaches.

Here, we revisit the taxonomy of the family *Rhizobiaceae* with a focus on the genus *Rhizobium*. To assist in refining the taxonomy of the genus *Rhizobium*, we performed whole genome sequencing of the type strains for 12 of the 13 *Rhizobium* species with validly published names and that previously lacked a publicly available genome sequence, and we generated new or more contiguous assembly for six additional *Rhizobiaceae* type strains. We then applied a previously established phylogenomic framework to reappraise the taxonomic classification of these 18 strains together with 224 additional *Rhizobiaceae* type strains with publicly available genome sequences. Based on the results of these analyses, we propose multiple taxonomic revisions to remove inconsistencies and ensure all validly named genera are monophyletic.

## MATERIALS AND METHODS

### Bacterial strains and growth conditions

All strains were grown using TY medium (5 g/L tryptone, 2.5 g/L yeast extract, 10 mM CaCl_2_, and 15 g/L agar for solid medium) at 28°C. *Rhizobium arsenicireducens* LMG 28795^T^, *Rhizobium cauense* LMG 26832^T^, *Rhizobium pakistanense* LMG 27895^T^, and *Rhizobium paranaense* LMG 27577^T^ were purchased from the BCCM/LMG strain collection (bccm.belspo.be). *Rhizobium capsici* JCM 19535^T^ and *Rhizobium straminoryzae* JCM 19536^T^ were purchased from the JCM strain collection (jcm.brc.riken.jp/en/).

*Allorhizobium paknamense* NBRC 109338^T^ and *Rhizobium puerariae* NBRC 110722^T^ were purchased from the NBRC strain collection (nite.go.jp/nbrc/catalogue/). *Rhizobium arsenicireducens* KCTC 72768^T^ and “*Affinirhizobium helianthi*” KCTC 23879^T^ was purchased from the KCTC strain collection (kctc.kribb.re.kr/en/). *Rhizobium acidisoli* LMG 18672^T^, *Rhizobium aegyptiacum* 1010^T^, *Rhizobium alamii* LMG 24466^T^, *Rhizobium azibense* HAMBI 3541^T^, *Rhizobium endophyticum* CCGE 2052^T^, *Rhizobium kunmingense* LMG 22609^T^, *Rhizobium mesosinicum* LMG 24135^T^, *Rhizobium soli* KCTC 12873^T^, *Rhizobium zeae* CRZMR18^T^, and *Endobacterium panacihumi* KCTC 62017^T^ were obtained from the strain collection maintained at the Universidad de Salamanca.

### 16S rRNA gene sequencing

Following receipt of the strains from the commercial strain collections, colony PCR and sequencing of the 16S rRNA genes of the strains was performed as described previously [8]. BLASTn, as implemented on the National Center for Biotechnology Information (NCBI) webserver, was used to compare the 16S rRNA genes to the NCBI core nucleotide database to confirm the identity of the strains prior to DNA isolation and whole genome sequencing. In addition, a local copy of BLASTn version 2.17.0+ [9] was used to compare the published 16S rRNA gene sequences of each type strain to the genome assemblies produced in the current study, to confirm the identity of each genome sequence.

### Whole genome sequencing

Single colonies were inoculated into TY broth and grown for one to two nights, following which DNA was extracted using either Monarch genomic DNA purification kits (New England Biolabs) or a DNeasy UltraClean Microbial Kit (Qiagen) according to the manufacturer’s instructions. Oxford Nanopore Technologies (ONT) library preparation was performed using either a Rapid Barcoding Kit 96 V14 (SQK-RBK114.96, ONT) or a Rapid PCR Barcoding Kit 24 V14 (SQK-RPB114.24, ONT) according to the manufacturer’s instructions, followed by sequencing on PromethION flow cell (R10.4.1, ONT) on a P2 Solo device. Basecalling and demultiplexing were performed using dorado version 0.7.4 with the model dna_r10.4.1_e8.2_400bps_sup@v4.3.0.

### Genome assembly and annotation

ONT reads were first filtered using Filtlong version 0.2.1 (github.com/rrwick/Filtlong). Genome assembly was then performed using the filtered ONT reads with Flye version 2.9.3 [10], and genome polishing performed using the filtered ONT reads and Medaka version 2.0.1 (github.com/nanoporetech/medaka). Next, FCS-adapter version 0.5.4 of the NCBI Foreign Contamination Screen (FCS) tool suite (github.com/ncbi/fcs) and custom code was used to identify, and if detected remove, adapter sequences from the polished assemblies, following which Pullseq version 1.0.2 (github.com/bcthomas/pullseq) was used to remove contigs less than 5000 bp in length.

Genome assembly quality was determined using CheckM version 1.2.3 [11] and initial taxonomic classification performed with the Genome Taxonomy Database Toolkit (GTDB-Tk) version 2.4.0 with database version R220 [12]. Genome annotation was then performed using the NCBI Prokaryotic Genome Annotation Pipeline (PGAP) version 2024-07-18.build7555 [13]. Accessions for genome assemblies generated as part of this study are provided in **Dataset S1**.

### Genome dataset

The core-proteome phylogeny and overall genomic relatedness indices (OGRIs) were calculated using the 18 genome assemblies produced as part of this study (**Dataset S1**) together with the genome assemblies of 227 additional type strains (**Datasets S2 and S3**). These 227 strains included type strains of all 219 *Rhizobiaceae* species for which we could find genomes available through the NCBI RefSeq database [14], an additional five *Rhizobiaceae* type strains of particular interest that were downloaded from the JGI Genome Portal [15], and three *Mesorhizobium* type strains available through NCBI RefSeq as an outgroup.

### Core-proteome phylogeny

An automated pipeline for constructing core-proteome phylogenies and calculating core-proteome average amino acid identity (cpAAI) values for the family *Rhizobiaceae* and order *Hyphomicrobiales* was prepared based on the framework and marker proteins described by Kuzmanović et al. [6] and diCenzo et al. [3]. This pipeline is freely available via GitHub (github.com/diCenzo-Lab/017_2025_Rhizobiaceae_taxonomy). For the current study, the pipeline was run in the *Rhizobiaceae* mode.

First, the 170 *Rhizobiaceae* marker genes of Kuzmanović et al. [6] were identified in the 245 genome sequences using tBLASTn version 2.17.0+ [9], aligned using MAFFT version 7.453 [16], trimmed using trimal version 1.4.rev22 with the automated1 algorithm [17], and finally concatenated. The concatenated alignment was then used to construct a maximum likelihood phylogeny using IQ-TREE2 version 2.2.2.4 [18]. First, ModelFinder [19], as implemented in IQ-TREE2, was used to identify the best scoring model based on Bayesian information criterion (BIC) with model search limited to LG models. Then, IQ-TREE2 was run using the best-scoring model (LG+F+I+R10) with branch support assessed using the Shimodaira–Hasegawa-like approximate likelihood ratio test (SH-aLRT) and ultrafast jackknife analysis with a subsampling proportion of 40%, with both metrics calculated from 1,000 replicates. Phylogenies were visualized using iTOL [20].

### Calculations of OGRIs

Average nucleotide identity (ANI) values were calculated using FastANI version 1.33 [21]. Digital DNA-DNA hybridization (dDDH) values were calculated using the Genome-to-Genome Distance Calculator version 3.0 (ggdc.dsmz.de/distcalc2.php) [22]. The cpAAI values were computed as described previously and were calculated as the proportion of differences in pairwise comparisons of the concatenated alignment whose construction is described in the previous section. Calculations were performed using the dist.aa() function of the ape package version 5.7.1 [23] in R version 4.4.1, with the scaled and pairwise.deletion options set to true.

## Data availability

Genome assemblies generated as part of this work, as well as the corresponding ONT reads, have been uploaded to the European Nucleotide Archive (ENA) under Project accession PRJEB102112; individual accessions for each genome and ONT read set are provided in **Dataset S1**. As the genomes had not completed processing at ENA by the time this manuscript was submitted, the fasta files were also added to our GitHub repository (github.com/diCenzo-Lab/017_2025_Rhizobiaceae_taxonomy).

The accessions for genome assemblies downloaded from NCBI RefSeq are provided in **Dataset S2**, while the accessions for the genome assemblies downloaded from the JGI Genome Portal are provided in **Dataset S3**. All code to repeat the analyses described in this study is available through GitHub (github.com/diCenzo-Lab/017_2025_Rhizobiaceae_taxonomy). The automated taxonomic pipeline is also available through the same GitHub repository to facilitate its reuse.

## RESULTS AND DISCUSSION

### An automated pipeline for taxonomic assignment

We previously published frameworks and computational workflows for the taxonomic assignment of species to genera in the family *Rhizobiaceae* [6], and genera to families in the order *Hyphomicrobiales* [3]. However, as these were multi-step workflows, they were not the easiest for others to apply to their own datasets or to troubleshoot. Here, we report an automated pipeline for cpAAI calculation and core-proteome phylogeny reconstruction, which is freely available for download through GitHub (github.com/diCenzo-Lab/017_2025_Rhizobiaceae_taxonomy). This single, one-line command uses the appropriate marker proteins described in our previous studies to produce a cpAAI matrix and a core-proteome maximum likelihood phylogeny. The pipeline can be run in either *Rhizobiaceae* mode to use the marker proteins of Kuzmanović et al. [6], or *Hyphomicrobiales* mode to use the marker proteins of diCenzo et al. [3].

Compared to our original pipeline, the primary change is the use of MAFFT [16] for marker protein alignment instead of Clustal Omega [24]. Although the marker protein sets were selected based on being present in all the strains included in the original studies, we found that with larger strain sets, one or more marker protein was occasionally absent, potentially due to the use of incomplete genome assemblies. For example, in the current study, 3 (1.2%) of the 245 genomes lacked between 1 and 2 of the 170 *Rhizobiaceae* marker genes. We were better able to handle these edge cases in our pipeline when switching to MAFFT, by adding “gap” proteins consisting solely of Xs prior to alignment; whereas MAFFT could handle these gap proteins, Clustal Omega could not. When comparing results for our current study to that of Kuzmanović et al. [6], we found that the cpAAI values of both studies were strongly correlated, although the values we calculated were on average 0.64% higher (standard deviation of 0.17) than those by Kuzmanović et al., with the difference decreasing as cpAAI values increased (**Figure S1**). We hypothesize that this difference is due to a combination of the use of a different aligner (MAFFT versus Clustal Omega) and the use of an expanded species dataset, which would result in slightly different protein alignments.

Kuzmanović et al. [6] proposed a cpAAI threshold of ∼ 86% for delineating genera. Here, we have adjusted this general threshold to ∼ 86.5% to account for the higher cpAAI calculated using our expanded dataset. However, as with Kuzmanović et al. [6], we consider this a guideline rather than a strict threshold, recognizing that higher or lower thresholds may be more appropriate for some genera to account for differences in evolution of each lineage.

### Whole genome sequencing of 18 *Rhizobiaceae* type strains

To facilitate the taxonomic refinement of the genus *Rhizobium*, we first generated whole genome sequences for 18 *Rhizobiaceae* type strains (**Table 1**). At the time of sequencing, 14 of these type strains had lacked a publicly available genome assembly, while the other four were available only as contigs and we aimed to generate more contiguous assemblies or complete genomes. Thirteen of the 18 genome assemblies are fully finished genomes (all replicons circularized); the remaining five have at least one replicon that was not circularized suggesting they are incomplete genomes although some of the non-circularized replicons could represent linear replicons [25]. Genome sizes ranged between 4.2 Mb and 7.5 Mb in size, with genome completeness scores of >99.4% and contamination scores <1.4% as determined by CheckM with the *Rhizobiaceae* marker gene set (**Table 1**). After accounting for the strains newly sequenced as part of this study, all but two *Rhizobium* species with validly published names now have a genuine genome sequence for their type strains, facilitating a more thorough taxonomic refinement of the genus *Rhizobium*. One of the species lacking an authentic genome for its type strain is *Rhizobium yanglingense*. Although a genome sequence is available for *R. yanglingense* LMG 19592^T^ through JGI (GOLD Project ID: Gp0220331), anomalies suggest that this genome sequence may not be of the true type strain and thus it was excluded from this study pending further investigation. The other *Rhizobium* species without a genome sequence for its type strain is *R. arsenicireducens*.

**Table 1.** Genome assembly statistics for type strains sequenced as part of this study.

| Strain | Assembly status | Genome size | Protein coding genes | Contigs | G+C Content (%) | Completeness | Contamination |
| --- | --- | --- | --- | --- | --- | --- | --- |
| *"Affinirhizobium helianthi"* KCTC 23879<sup>T</sup> | Draft | 4,252,791 | 3,820 | 6 | 57.86 | 99.96 | 0.00 |
| *Allorhizobium paknamense* NBRC 109338<sup>T</sup> | Draft | 5,413,673 | 4,728 | 10 | 59.84 | 99.93 | 0.59 |
| *Endobacterium panacihumi* KCTC 62017<sup>T</sup> | Complete | 5,901,858 | 5,565 | 7 | 59.29 | 99.81 | 1.23 |
| *Rhizobium acidisoli* LMG 28672<sup>T</sup> | Complete | 6,940,437 | 6,459 | 4 | 61.31 | 100 | 0.67 |
| *Rhizobium aegyptiacum* 1010<sup>T</sup> | Complete | 6,368,843 | 5,945 | 6 | 61.23 | 100 | 0.30 |
| *Rhizobium alamii* LMG 24466<sup>T</sup> | Complete | 7,434,066 | 7,099 | 3 | 60.01 | 100 | 1.05 |
| *Rhizobium azibense* HAMBI 3541<sup>T</sup> | Complete | 7,092,048 | 6,454 | 4 | 59.55 | 100 | 0.48 |
| *Rhizobium capsici* JCM 19535<sup>T</sup> | Draft | 4,526,232 | 4,144 | 9 | 55.95 | 99.93 | 0.11 |
| *Rhizobium cauense* LMG 26832<sup>T</sup> | Complete | 6,938,506 | 6,327 | 3 | 59.13 | 100 | 1.33 |
| *Rhizobium endophyticum* CCGE 2052<sup>T</sup> | Complete | 6,156,979 | 5,878 | 3 | 61.22 | 100 | 0.00 |
| *Rhizobium kunmingense* LMG 22609<sup>T</sup> | Draft | 5,510,491 | 5,159 | 4 | 61.07 | 99.96 | 1.00 |
| *Rhizobium mesosinicum* LMG 24135<sup>T</sup> | Complete | 6,131,211 | 5,812 | 3 | 59.94 | 100 | 0.44 |
| *Rhizobium pakistanense* LMG 27895<sup>T</sup> | Complete | 5,476,458 | 5,215 | 2 | 61.94 | 100 | 0.04 |
| *Rhizobium paranaense* LMG 27577<sup>T</sup> | Complete | 6,861,319 | 6,351 | 4 | 59.56 | 99.98 | 0.33 |
| *Rhizobium puerariae* NBRC 110722<sup>T</sup> | Complete | 6,632,821 | 6,262 | 4 | 62.66 | 100 | 0.00 |
| *Rhizobium soli* KCTC 12873<sup>T</sup> | Draft | 5,187,886 | 4,820 | 6 | 60.06 | 99.48 | 0.09 |
| *Rhizobium straminoryzae* JCM 19536<sup>T</sup> | Complete | 5,258,508 | 4,747 | 4 | 63.69 | 100 | 1.33 |
| *Rhizobium zeae* CRZMR18<sup>T</sup> | Complete | 6,273,448 | 5,859 | 3 | 59.39 | 99.52 | 0.37 |

### Taxonomic status of *R. arsenicireducens*

As part of this study, we attempted to sequence *R. arsenicireducens* LMG 28795^T^; however, 16S rRNA gene sequencing indicated that the strain received from BCCM/LMG was *Escherichia coli*, which was further confirmed based on whole genome sequencing and taxonomic classification with the GTDB-Tk. BCCM/LMG subsequently confirmed that they were only able to isolate *E. coli* from their stocks of strain LMG 28795^T^ and thus this strain was removed from their collection. Similarly, 16S rRNA gene sequencing of *R. arsenicireducens* KCTC 72768^T^ failed to identify a *Rhizobium* strain. As we were unable to obtain *R. arsenicireducens* MTCC 12115^T^, we were unable to confirm the identity of this strain. We suggest that the identity of *R. arsenicireducens* MTCC 12115^T^ be confirmed by those with access to the strain, and if this strain has also been lost, that a neotype should be designated according to Rule 18c, or otherwise that the taxonomic status of *R. arsenicireducens* should be revised.

### Identification of synonymous species names

ANI (as determined by FastANI) and dDDH were used to evaluate whether any of the species included in our study are synonymous. Ten pairs of type strains had ANI values ≥95%, eight of which were 96.02% or lower (**Table 2**). As the dDDH values for all eight pairs of type strains with ANI values between 95% and 96.02% were below 70%, and considering that an ANI threshold of 96% has been proposed as an appropriate cut-off for species delineation in the *Rhizobium* species complex [26], we consider these eight pairs of species to be non-synonymous. On the other hand, the ANI and dDDH values for the comparison of *R. aegyptiacum* 1010^T^ and *Rhizobium aethiopicum* HBR26^T^ were 98.4% and 86.6%, respectively, and thus we propose that *R. aegyptiacum* is a later heterotypic synonym of *R. aethiopicum*. Similarly, the ANI and dDDH values for the comparison of “*Shinella sumterensis*” MEC087^T^ and “*Shinella oryzae*” Z-25^T^ were 96.5% and 72%, respectively, leading us to suggest that these two names are synonymous although neither has been validly published.

**Table 2.**
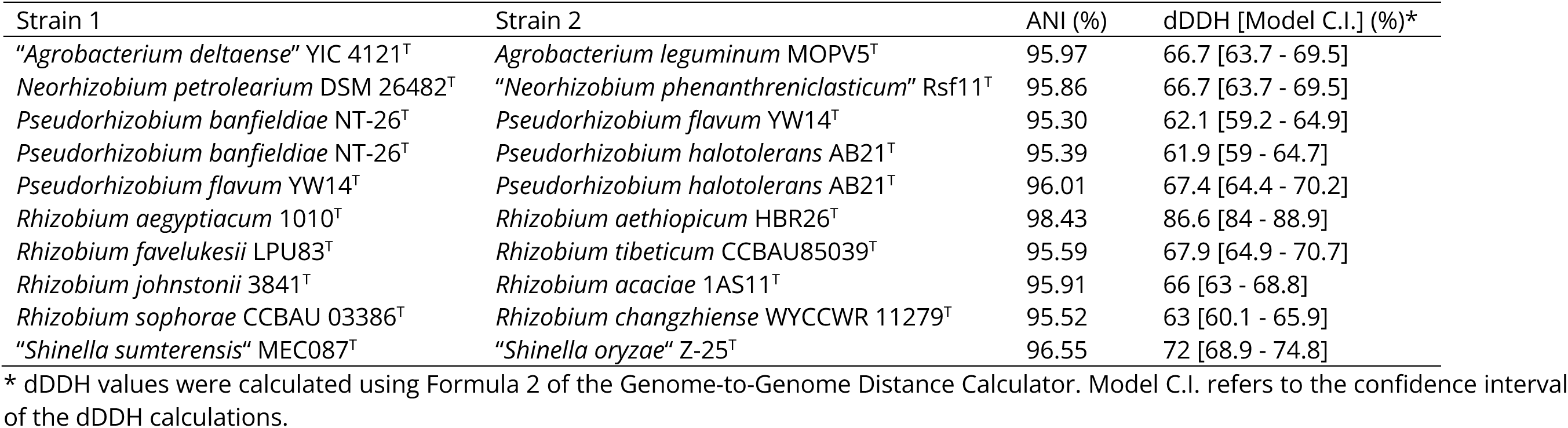
Average nucleotide identity (ANI) and digital DNA-DNA hybridization (dDDH) for species pairs with ANI values ≥ 95%.

| Strain 1 | Strain 2 | ANI (%) | dDDH [Model C.I.] (%)<sup>*</sup> |
| --- | --- | --- | --- |
| *"Agrobacterium deltaense"* YIC 4121<sup>T</sup> | *Agrobacterium leguminum* MOPV5<sup>T</sup> | 95.97 | 66.7 [63.7 - 69.5] |
| *Neorhizobium petrolearium* DSM 26482<sup>T</sup> | *"Neorhizobium phenanthreniclasticum"* Rsf11<sup>T</sup> | 95.86 | 66.7 [63.7 - 69.5] |
| *Pseudorhizobium banfieldiae* NT-26<sup>T</sup> | *Pseudorhizobium flavum* YW14<sup>T</sup> | 95.30 | 62.1 [59.2 - 64.9] |
| *Pseudorhizobium banfieldiae* NT-26<sup>T</sup> | *Pseudorhizobium halotolerans* AB21<sup>T</sup> | 95.39 | 61.9 [59 - 64.7] |
| *Pseudorhizobium flavum* YW14<sup>T</sup> | *Pseudorhizobium halotolerans* AB21<sup>T</sup> | 96.01 | 67.4 [64.4 - 70.2] |
| *Rhizobium aegyptiacum* 1010<sup>T</sup> | *Rhizobium aethiopicum* HBR26<sup>T</sup> | 98.43 | 86.6 [84 - 88.9] |
| *Rhizobium favelukesii* LPU83<sup>T</sup> | *Rhizobium tibeticum* CCBAU85039<sup>T</sup> | 95.59 | 67.9 [64.9 - 70.7] |
| *Rhizobium johnstonii* 3841<sup>T</sup> | *Rhizobium acaciae* 1AS11<sup>T</sup> | 95.91 | 66 [63 - 68.8] |
| *Rhizobium sophorae* CCBAU 03386<sup>T</sup> | *Rhizobium changzhiense* WYCCWR 11279<sup>T</sup> | 95.52 | 63 [60.1 - 65.9] |
| *"Shinella sumterensis"* MEC087<sup>T</sup> | *"Shinella oryzae"* Z-25<sup>T</sup> | 96.55 | 72 [68.9 - 74.8] |

<sup>\*</sup> dDDH values were calculated using Formula 2 of the Genome-to-Genome Distance Calculator. Model C.I. refers to the confidence interval of the dDDH calculations.

### Rhizobium azibense and Rhizobium gallicum are not synonymous

Our data are inconsistent with those of Volpiano et al. [27], who recently proposed that *R. azibense* is a later heterotypic synonym of *R. gallicum* based on the ANI value between *R. azibense* 23C2^T^ and *R. gallicum* SEMIA 4080 being >99%. Volpiano et al. also noted that the 16S rRNA gene sequence of the *R. azibense* 23C2^T^ genome sequence was 99.13% identical to the sequence reported in the original genome description by Mnasri et al. [28]. Here, we generated a genome sequence for *R. azibense* HAMBI 3541^T^. Comparison of the 16S rRNA gene sequence of this genome assembly to that reported by Mnasri et al. [28] revealed a 100% match, confirming the genome represents the authentic type strain. In addition, the ANI value between *R. azibense* HAMBI 3541^T^ and *R. azibense* 23C2^T^ was only 92.6%, suggesting that the genome sequence generated by Volpiano et al. [27] does not represent the authentic *R. azibense* type strain. When comparing our genome sequence for *R. azibense* 23C2^T^ with the published genome sequence for *R. gallicum* R602^T^, the ANI value was 92.5%, consistent with *R. azibense* and *R. gallicum* representing distinct species rather than being synonymous.

### Genus-level refinements to the taxonomy of the family *Rhizobiaceae*

Using our taxonomic pipeline, we constructed a maximum likelihood phylogeny of the 242 *Rhizobiaceae* type strains and three *Mesorhizobium* type strains as an outgroup (**Figure S2**) and calculated cpAAI values between each pair of species (**Dataset S4**). These data were used to reassign species to different genera, as appropriate, to ensure each genus is monophyletic. In cases where it was unclear whether a species should be transferred to an existing or novel genus, cpAAI was employed, generally using a threshold of ∼ 86.5% to determine genus affiliation. At the same time, we generally chose to limit the number of taxonomic revisions required to ensure monophyly of each genus. In other words, we generally avoid making proposal to split a monophyletic genus based solely on cpAAI unless the evidence to do so is clear.

Proposals to reclassify Hoeflea poritis as Allohoeflea poritis gen. nov. comb. nov., Pseudohoeflea coraliihabitans as Neohoeflea coraliihabitans gen. nov. comb. nov., and to transfer Hoeflea alexandrii, Hoeflea halophila, Hoeflea olei, and Hoeflea phototrophica to Parahoeflea gen. nov.

In our phylogeny, the genus *Hoeflea* was polyphyletic (**Figure 1**). Of the nine species assigned to the genus *Hoeflea*, seven grouped with the type species *Hoeflea marina*, while *H. poritis* E7-10^T^ and “*H. prorocentri*” PM5-8^T^ formed a distinct monophyletic group as sister taxa to the genus *Flavimaribacter*. The cpAAI value between *H. poritis* E7-10^T^ and “*H. prorocentri*” PM5-8^T^ was ∼ 85.6%, which is reasonably close to the proposed genus delimitation threshold of ∼ 86.5%, and thus we propose that they belong to the same genus. In addition, the cpAAI values calculated between *Flavimaribacter sediminis* WL0058^T^ and either *H. poritis* E7-10^T^ or “*H. prorocentri*” PM5-8^T^ were less than 78%, suggesting that these latter two species belong to a novel genus distinct from *Flavimaribacter*. We therefore suggest that *H. poritis* and “*H. prorocentri*” belong to the same genus and be transferred to *Allohoeflea* gen. nov.. A formal description of *Allohoeflea poritis* comb. nov. is provided below; however, as “*H. prorocentri*” is not yet validly published, reclassification of this species currently cannot be formally proposed.

**Figure 1.**
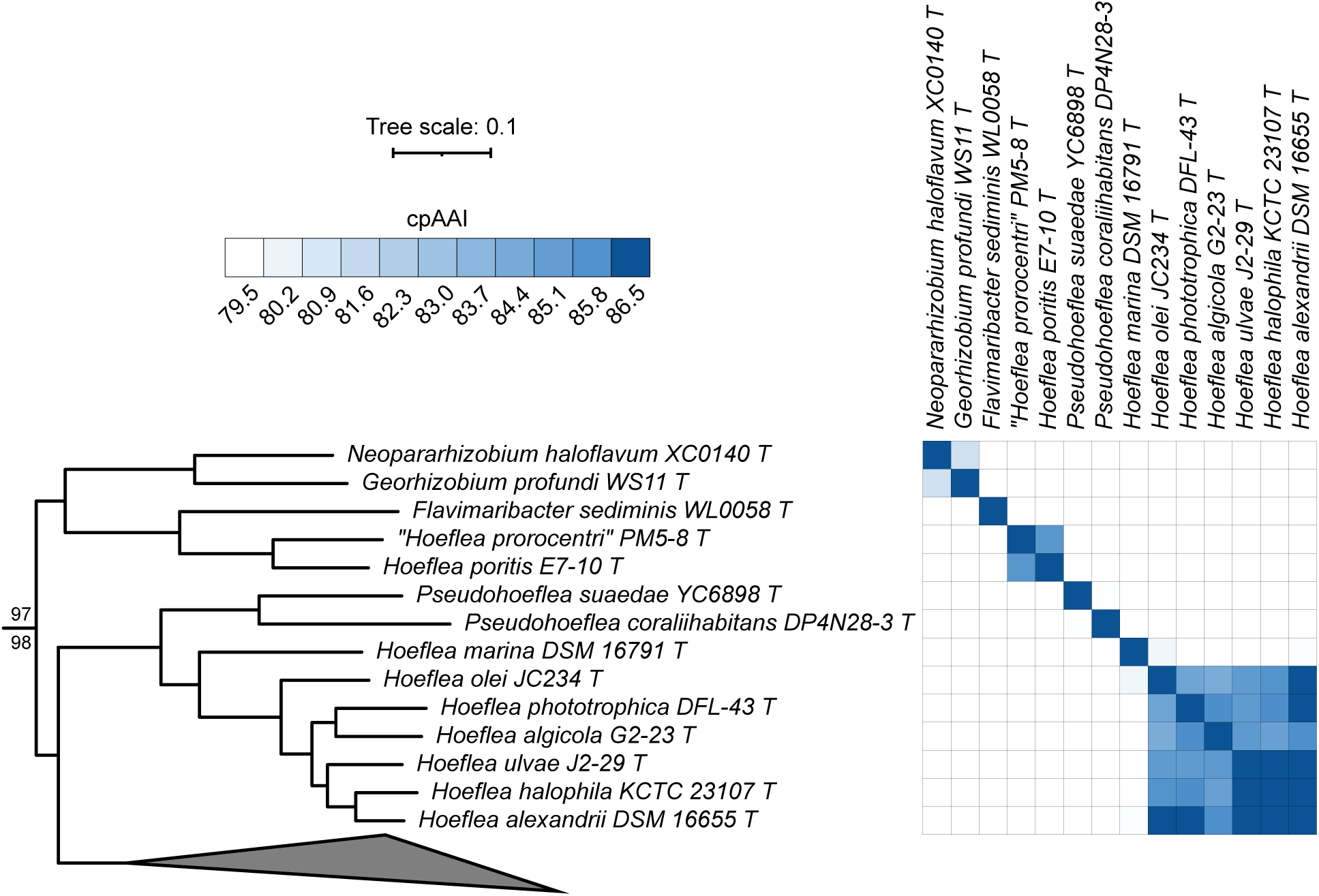
Phylogenomic analysis of the genera *Hoeflea* and *Pseudohoeflea*. (**Left**) A maximum likelihood phylogeny of the genera *Hoeflea, Pseudohoeflea*, *Flavimaribacter*, *Georhizobium*, and “*Neopararhizobium*” extracted from the full *Rhizobiaceae* phylogeny of Figure S2. The numbers on the branches indicate support values by SH-aLRT (above) and ultra-fast jackknife (below); values are only shown at nodes where at least one value is below 100. The scale bar represents the average number of amino acid substitutions per site. (**Right**) A matrix showing the core-proteome average amino acid identity (cpAAI) values between each pair of strains calculated using a core set of 170 proteins. Values less than 79.5% are in white while all values greater than 86.5% are in the same shade of blue. An interactive version of this phylogeny is available at itol.embl.de/shared/1ps8ayxRcrNDU.

The remaining seven *Hoeflea* species formed a monophyletic group, of which the deepest branching strain was the type strain of the genus type species, *H. marina* DSM 16791^T^ (**Figure 1**). However, the cpAAI values between *H. marina* DSM 16791^T^ and the other six *Hoeflea* type strains were all < 80.1% (**Figure 1**). This suggests that *H. marina* is the only species belonging to the genus *Hoeflea*, and that the other six species should be reclassified to a novel genus. Arguably, the cpAAI values between these six species would support their reclassification into three novel genera (**Figure 1**). However, as all pairwise cpAAI values were > 84% (and most were over 85%) and thus reasonably close to the proposed genus delimitation threshold of ∼ 86.5% without a clean separation of the three sub-clades, and because the six species formed a monophyletic group (**Figure 1**), we instead suggest that all six species be transferred to a single genus, *Parahoeflea* gen. nov.. Formal descriptions are provided below for those species (*Parahoeflea alexandrii* comb. nov., *Parahoeflea halophila* comb. nov., *Parahoeflea olei* comb. nov., and *Parahoeflea phototrophica* comb. nov.) whose type strains currently meet the availability criteria specified by the ICNP.

Lastly, although *Pseudohoeflea suaedae* YC6898^T^ and *P. coraliihabitans* DP4N28-3^T^ formed a monophyletic group, the pairwise cpAAI value between these two strains was < 80% (**Figure 1**). Given that this cpAAI value is much lower that the proposed genus delimitation threshold of ∼ 86.5%, we argue that these two species belong to distinct genera and therefore propose that *P. coraliihabitans* be reclassified as *Neohoeflea coraliihabitans* gen. nov. comb. nov., whose formal description is provided below.

Proposals to reclassify Rhizobium aquaticum as Limnomicrobium aquaticum gen. nov. comb. nov., Rhizobium alvei as Fluviimicrobium alvei gen. nov., comb. nov., and Rhizobium setariae as Yannia setariae gen. nov. comb. nov.

*R. aquaticum* DSM 29780^T^, *R. alvei* TNR-22^T^, and *R. setariae* KVB221^T^ did not group with the genus *Rhizobium* but instead formed a monophyletic group with *Onobrychidicola muellerharveyae* TN2^T^, the type strain of the type species of the genus *Onobrychidicola* (**Figure 2**). The pairwise cpAAI values between *R. aquaticum* DSM 29780^T^ and the other three strains in this clade were all < 75.2% (**Figure 2**), suggesting this species belongs to its own genus and leading us to propose that *R. aquaticum* be reclassified as *Limnomicrobium aquaticum* gen. nov. comb. nov., whose formal description is provided below. In addition, the pairwise cpAAI values of *O. muellerharveyae* TN2^T^ with *R. alvei* TNR-22^T^ and *R. setariae* KVB221^T^ were ∼ 84.0% and ∼ 81.9%, respectively, while the cpAAI value between *R. alvei* TNR-22^T^ and *R. setariae* KVB221^T^ was ∼ 82.5% (**Figure 2**). These results suggest that *O. muellerharveyae* TN2^T^, *R. alvei* TNR-22^T^, and *R. setariae* KVB221^T^ each belong to a distinct genus. As a result, we propose that *R. alvei* be reclassified as *Fluviimicrobium alvei* gen. nov., comb. nov., and that *R. setariae* be reclassified as *Yannia setariae* gen. nov. comb. nov., whose formal descriptions are provided below.

**Figure 2.**
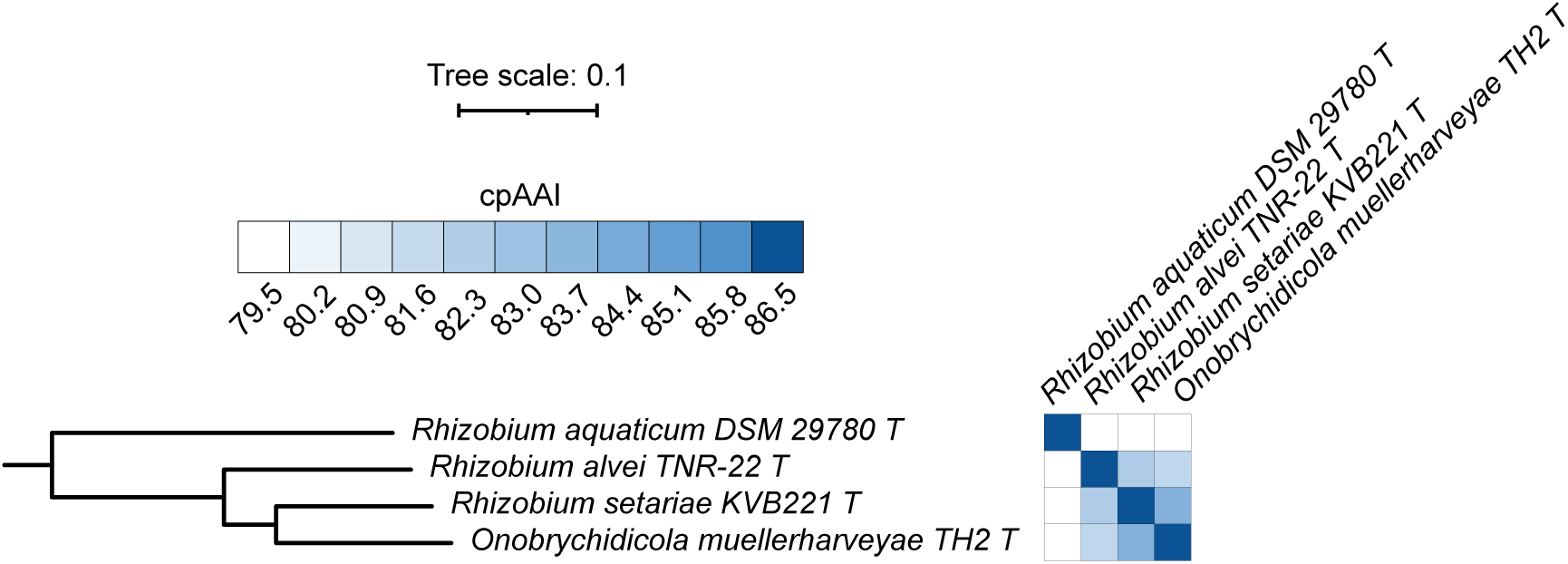
Phylogenomic analysis of the genus *Onobrychidicola* and related species. (**Left**) A maximum likelihood phylogeny of the genus *Onobrychidicola* and related species is provided; this phylogeny was extracted from the full *Rhizobiaceae* phylogeny of Figure S2. All support values calculated by SH-aLRT and ultra-fast jackknife were 100 and are therefore not shown. The scale bar represents the average number of amino acid substitutions per site. (**Right**) A matrix showing the core-proteome average amino acid identity (cpAAI) values between each pair of strains calculated using a core set of 170 proteins. Values less than 79.5% are in white while all values greater than 86.5% are in the same shade of blue. An interactive version of this phylogeny is available at itol.embl.de/shared/1ps8ayxRcrNDU.

Proposal to reclassify *Ensifer aridi* as *Sinorhizobium aridi* comb. nov.

*E. aridi* LMR001^T^ fell within the genus *Sinorhizobium* in our maximum likelihood phylogeny (**Figure S2**) and shared higher cpAAI values with the *Sinorhizobium* type strains (∼ 91.3% to ∼ 93.5%) than with the *Ensifer* type strains (∼ 88.9% to ∼ 89.4%) (**Dataset S4**). In particular, the cpAAI values for the type strains of the type species are 92.27% for *Sinorhizobium fredii* and 89.06% for *Ensifer adhaerens*. We therefore propose that *E. aridi* be reclassified as *Sinorhizobium aridi* comb. nov., whose formal description is provided below.

### Proposals to transfer the species of the “tropici-rhizogenes” and “tubonense-tumorigenes” clades of the genus *Rhizobium* to *Martinezia* gen. nov. and *Arminia* gen. nov., respectively

The genus *Rhizobium* can be broadly divided into three sub-clades [29], the “leguminosarum-etli” clade that includes the type species of the genus *Rhizobium*, the “tropici-rhizogenes” clade, and the “tubonense-tumorigenes” clade (**Figure 3**). To date, two of the three known species of the “tubonense-tumorigenes” clade are plant tumorigenic bacteria [29–31], whereas the other two clades are dominated by rhizobia. Most of the within-clade cpAAI comparisons are > 88% while nearly all between-clade cpAAI comparisons are < 86.5% (**Figure 3**). The exception is *Rhizobium tubonense* CCBAU 85046^T^, which displays pairwise cpAAI values from ∼ 87.1% and ∼87.9% with all species of both the “tropici-rhizogenes” and “tubonense-tumorigenes” clades (**Figure 3**). In addition, the genus “*Oryzifoliimicrobium*” was recently proposed to encompass the species “*Oryzifoliimicrobium ureilyticum*” [32]. In our analysis, “*O. ureilyticum*” SG148^T^ falls between the “leguminosarum-etli” and “tropici-rhizogenes” clades within the genus *Rhizobium*, which would result in the genus *Rhizobium* being paraphyletic. Moreover, “*O. ureilyticum*” SG148^T^ shares < 84% cpAAI with all but one *Rhizobium* type strains, consistent with the genera *Rhizobium* and “*Oryzifoliimicrobium*” not being synonymous. Therefore, based on the weight of the evidence, we propose that the genus *Rhizobium* be split into three genera, with the genus *Rhizobium* encompassing the “leguminosarum-etli” clade, *Martinezia* gen. nov. encompassing the “tropici-rhizogenes” clade, and *Arminia* gen. nov. encompassing the “tubonense-tumorigenes” clade. While we note that cpAAI alone was not sufficient to determine if *R. tubonense* CCBAU 85046^T^ belongs to the genus *Martinezia* gen. nov. or the genus *Arminia* gen. nov., considering the location of this strain in the phylogeny (**Figure 3**) and that the cpAAI values between this strain and *Arminia tumorigenes* 1078^T^ comb. nov. and *Arminia rhododendri* rho-6.2^T^ comb. nov. are ∼ 87.2%, we propose that *R. tubonense* should be transferred to *Arminia* gen. nov.. Formal descriptions are provided below for *Martinezia* gen. nov., *Arminia* gen. nov., and for those species (*Arminia rhododendri* comb. nov., *Arminia tubonensis* comb. nov., *Arminia tumorigenes* comb. nov., *Martinezia calliandrae* comb. nov., *Martinezia dioscoreae* comb. nov., *Martinezia freirei* comb. nov., *Martinezia hainanensis* comb. nov., *Martinezia jaguaris* comb. nov., *Martinezia leucaenae* comb. nov., *Martinezia lusitana* comb. nov., *Martinezia multihospitum* comb. nov., *Martinezia miluonensis* comb. nov., *Martinezia paranaensis* comb. nov., *Martinezia rhizogenes* comb. nov., and *Martinezia tropici* comb. nov.) whose type strains currently meet the availability criteria specified by the ICNP.

**Figure 3.**
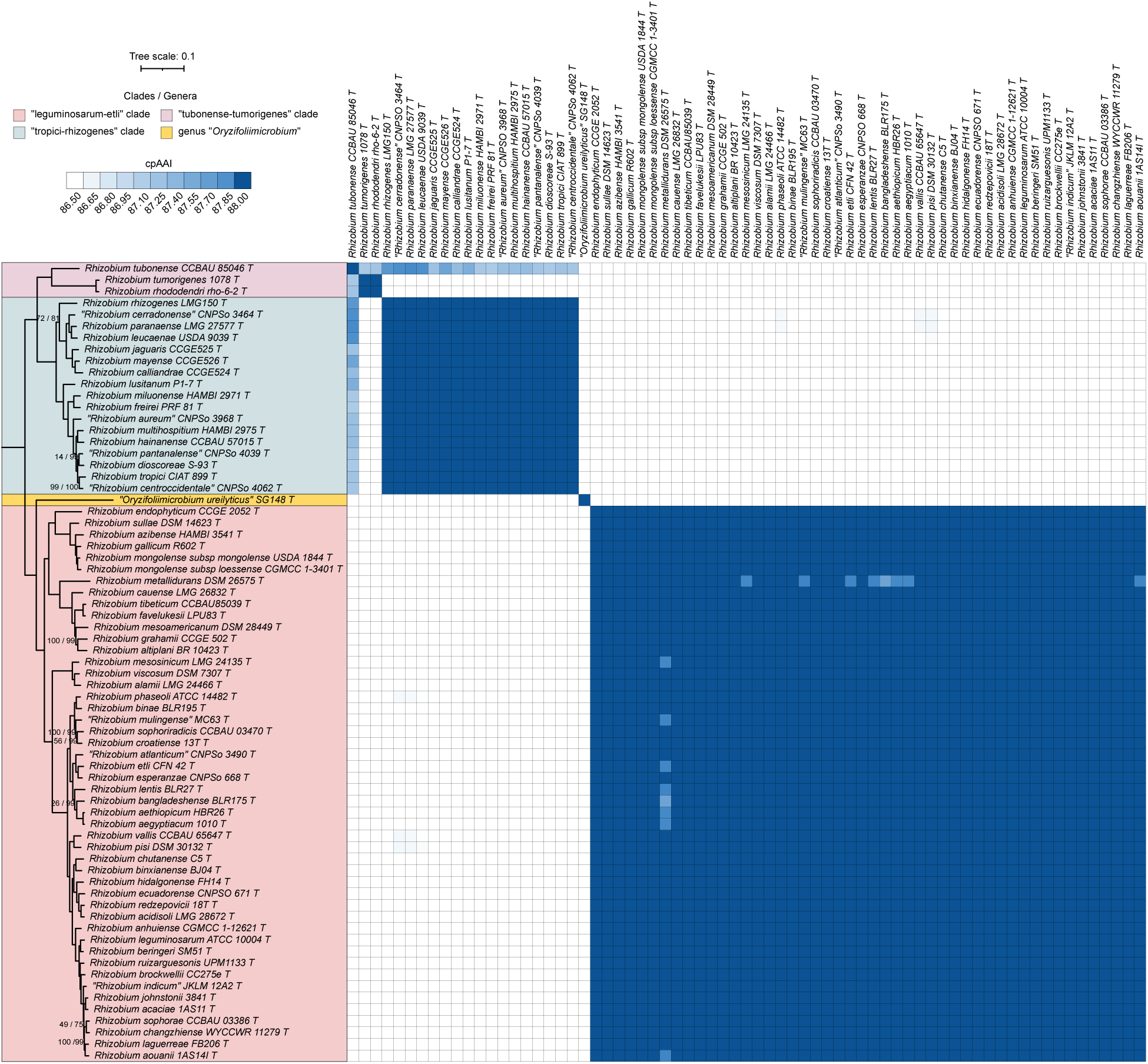
Phylogenomic analysis of the genera *Rhizobium* and “*Oryzifoliimicrobium*”. (**Left**) A maximum likelihood phylogeny of the genera *Rhizobium* and “*Oryzifoliimicrobium*” is provided; this phylogeny was extracted from the full *Rhizobiaceae* phylogeny of Figure S2. The numbers on the branches indicate support values by SH-aLRT (above) and ultra-fast jackknife (below); values are only shown at nodes where at least one value is below 100. The scale bar represents the average number of amino acid substitutions per site. (**Right**) A matrix showing the core-proteome average amino acid identity (cpAAI) values between each pair of strains calculated using a core set of 170 proteins. Values less than 86.5% are in white while all values greater than 88.0% are in the same shade of blue. An interactive version of this phylogeny is available at itol.embl.de/shared/1ps8ayxRcrNDU.

Proposal to transfer Allorhizobium ampelinum, Allorhizobium taibaishanense, Allorhizobium terrae, and Allorhizobium vitis to Gillisella gen. nov.

Although the genus *Allorhizobium* is monophyletic, when compared to *Allorhizobium undicola* ATCC 700741^T^, the type strain of the type species of the genus *Allorhizobium*, *A. paknamense* NBRC 109338^T^ was the only *Allorhizobium* type strain to display a cpAAI value > 86.5%; all others were below 85% (**Figure 4**). *A. sonneratiae* BGMRC 0089^T^ forms its own lineage as a sister taxon to *A. undicola* ATCC 700741^T^ and *A. paknamense* NBRC 109338^T^ (**Figure 4**). The pairwise cpAAI values between *A. sonneratiae* BGMRC 0089^T^ and *A. undicola* ATCC 700741^T^ or *A. paknamense* NBRC 109338^T^ is 84.0% and 84.9%, respectively, suggesting that *A. sonneratiae* BGMRC 0089^T^ belongs to a novel genus (**Figure 4**). However, as the *A. sonneratiae* type strain currently does not meet the availability criteria specified by the ICNP, a new combination for this species currently cannot be validly proposed.

**Figure 4.**
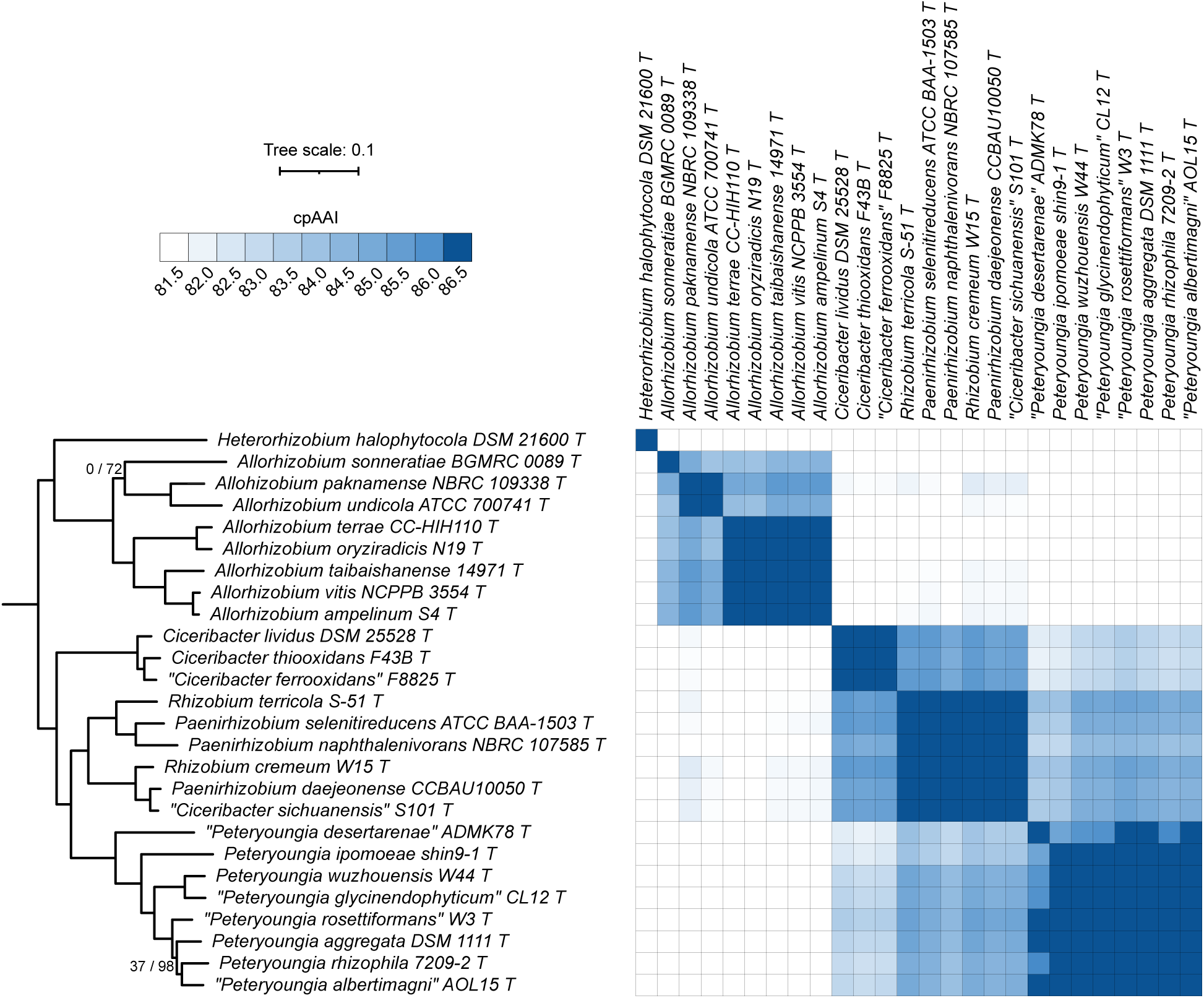
Phylogenomic analysis of the genera *Allorhizobium* and *Paenirhizobium*. (**Left**) A maximum likelihood phylogeny of the genera *Allorhizobium, Ciceribacter*, *Paenirhizobium*, and *Peteryoungia*, together with related species is provided; this phylogeny was extracted from the full *Rhizobiaceae* phylogeny of Figure S2. The numbers on the branches indicate support values by SH-aLRT (left) and ultra-fast jackknife (right); values are only shown at nodes where at least one value is below 100. The scale bar represents the average number of amino acid substitutions per site. (**Right**) A matrix showing the core proteome average amino acid identity (cpAAI) values between each pair of strains calculated using a core set of 170 proteins. Values less than 81.5% are in white while all values greater than 86.5% are in the same shade of blue. An interactive version of this phylogeny is available at itol.embl.de/shared/1ps8ayxRcrNDU.

*Allorhizobium terrae* CC-HIH110^T^, *A. oryziradicis* N19^T^, *A. taibaishanense* 14971^T^, *A. vitis* NCPPB 3554^T^, and *A. ampelinum* S4^T^ formed a monophyletic group in our phylogeny (**Figure 4**). The cpAAI values between these five strains were all > 86.5% (and most were above 87.0%), while the values between these five strains and the other three members of this clade were between 84% and 85.7% (with most below 85%) (**Figure 4**). Consequently, we suggest that these five species belong to a common genus that is separate from the other three species of this clade. We therefore propose that these five species be transferred to the genus *Gillisella* gen. nov.. Formal descriptions are provided below for those species (*Gillisella ampelina* comb. nov., *Gillisella taibaishanensis* comb. nov., *Gillisella terrae* comb. nov., and *Gillisella vitis* comb. nov.) whose type strains currently meet the availability criteria specified by the ICNP.

### Proposal to reclassify Rhizobium cremeum as Paenirhizobium cremeum comb. nov

*R. terricola* S-51^T^ and *R. cremeum* W15^T^ did not group with the genus *Rhizobium* but instead clustered with the three type strains of the genus *Paenirhizobium* (**Figure 4**). Likewise, “*Ciceribacter sichuanensis*” S101^T^ clusters with the type strains of the genus *Paenirhizobium* (**Figure 4**). Additionally, the genus *Paenirhizobium* was not monophyletic without the inclusion of these three type strains, and all pairwise cpAAI values between the six strains were > 86.5% with most > 87.5% (**Figure 4**). Collectively, these data suggest that *R. terricola*, *R. cremeum*, and “*C. sichuanensis*” should be transferred to the genus *Paenirhizobium*. We therefore propose that *R. cremeum* be reclassified as *Paenirhizobium cremeum* comb. nov., whose formal description is provided below. As “*C. sichuanensis*” is not yet validated and as *R. terricola* S-51^T^ currently does not meet the availability criteria specified by the ICNP, we are currently unable to formally propose new combinations for these species.

### Proposal to transfer Rhizobium capsici and Rhizobium straminoryzae to the genus Affinirhizobium

*R. oryzicola* 5753^T^, *R. straminoryzae* JCM 19536^T^, and *R. capsici* JCM 19535^T^ did not group with the genus *Rhizobium* but instead clustered with the five type strains of the genus *Affinirhizobium* (**Figure 5**). Additionally, the genus *Affinirhizobium* was not monophyletic without the inclusion of *R. straminoryzae* JCM 19536^T^ and *R. capsici* JCM 19535^T^. Excluding *R. oryzicola* 5753^T^ and "*Affinirhizobium gouqiense*” SSA5-23^T^, the pairwise cpAAI values between each of the remaining six type strains were all above 86.5% (**Figure 5**). On the other hand, the pairwise cpAAI values were above 86.5% for only two of the seven comparisons between *R. oryzicola* 5753^T^ or "*A. gouqiense*” SSA5-23^T^ and the other members of this clade (**Figure 5**). Nevertheless, as all pairwise cpAAI values were > 85.7% and thus reasonably close to the proposed genus delimitation threshold of ∼ 86.5% without a clean separation of the species, and because the eight species formed a monophyletic group (**Figure 5**), we argue that all eight species belong to a single genus. Consequently, we propose that *R. capsici* be reclassified as *Affinirhizobium capsici* comb. nov. and that *R. straminoryzae* be reclassified as *Affinirhizobium straminoryzae* comb. nov., whose formal descriptions are provided below. As the *R. oryzicola* type strain currently does not meet the availability criteria specified by the ICNP, a new combination for this species currently cannot be validly proposed.

**Figure 5.**
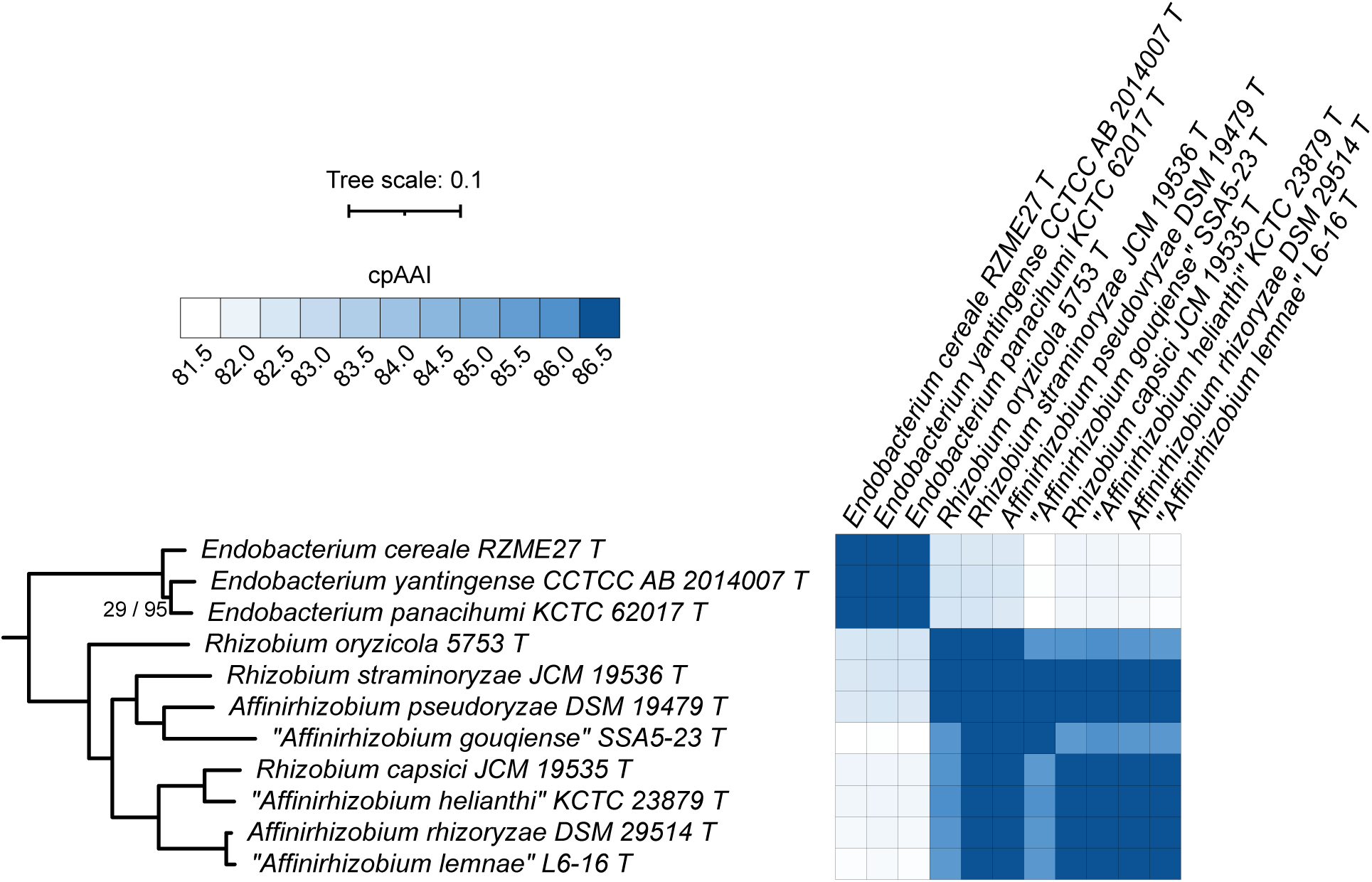
Phylogenomic analysis of the genus *Affinirhizobium*. (**Left**) A maximum likelihood phylogeny of the genera *Endobacterium*, and *Affinirhizobium*; this phylogeny was extracted from the full *Rhizobiaceae* phylogeny of Figure S2. The numbers on the branches indicate support values by SH-aLRT (left) and ultra-fast jackknife (right); values are only shown at nodes where at least one value is below 100. The scale bar represents the average number of amino acid substitutions per site. (**Right**) A matrix showing the core proteome average amino acid identity (cpAAI) values between each pair of strains calculated using a core set of 170 proteins. Values less than 81.5% are in white while all values greater than 86.5% are in the same shade of blue. An interactive version of this phylogeny is available at itol.embl.de/shared/1ps8ayxRcrNDU.

Proposals to reclassify Rhizobium puerariae as Neorhizobium puerariae comb. nov., Rhizobium zeae as Aliirhizobium zeae comb. nov., and Rhizobium soli as Velazquezia soli gen. nov. comb. nov. The *Neorhizobium* – *Aliirhizobium* – *Terrirhizobium* clade of the family *Rhizobiaceae* consists of 22 species, including five currently assigned to the genus *Rhizobium* (**Figure 6**). This clade is not fully resolved at a cpAAI threshold of 86.5%; however, clear genus boundaries emerge at a cpAAI threshold of 88.0% to 88.5% that align with the phylogeny and that allow for the separation of all three currently named genera. The pairwise cpAAI values between the 12 type strains that form a clade (*Neorhizobium sensu stricto*) that includes *Neorhizobium galegae* range from 88.9% to 96.9%, while the pairwise cpAAI values between the 12 type strains of *Neorhizobium sensu stricto* and the other 10 type strains range from 80.2% to 88.3% (**Figure 6**). Likewise, the pairwise cpAAI values for the five type strains that include all named *Aliirhizobium* species range from 90.7% to 94.1%, while the pairwise cpAAI values between those five type strains and the remaining 17 type strains range from 84.2% to 86.5% (**Figure 6**). The cpAAI values between *Terrirhizobium terrae* CC-CFT758^T^, the sole member of the genus *Terrirhizobium*, and the other 21 type strains range from 79.2% to 84.4% (**Figure 6**). *Rhizobium soli* KCTC 12873^T^ forms its own lineage that is sister to *Aliirhizobium* and *Terrirhizobium*, and shares cpAAI values of between 80.7% and 86.7% with the other 21 species of this clade (**Figure 6**). Likewise, *Rhizobium pakistanense* LMG 27895^T^ forms its own lineage as an outgroup to the rest of *Neorhizobium* – *Aliirhizobium* – *Terrirhizobium* clade, and shares cpAAI values of between 79.4% and 87.9% with the other 21 species of this clade (**Figure 6**). Lastly, the pairwise cpAAI value between the sister taxa “*Neorhizobium lilium*” 24NR^T^ and “*Neorhizobium deserti*” SPY-1^T^ is 88.3%, while the cpAAI values between these strains and the other 20 type strains of this clade range from 80.0% to 88.3%.

**Figure 6.**
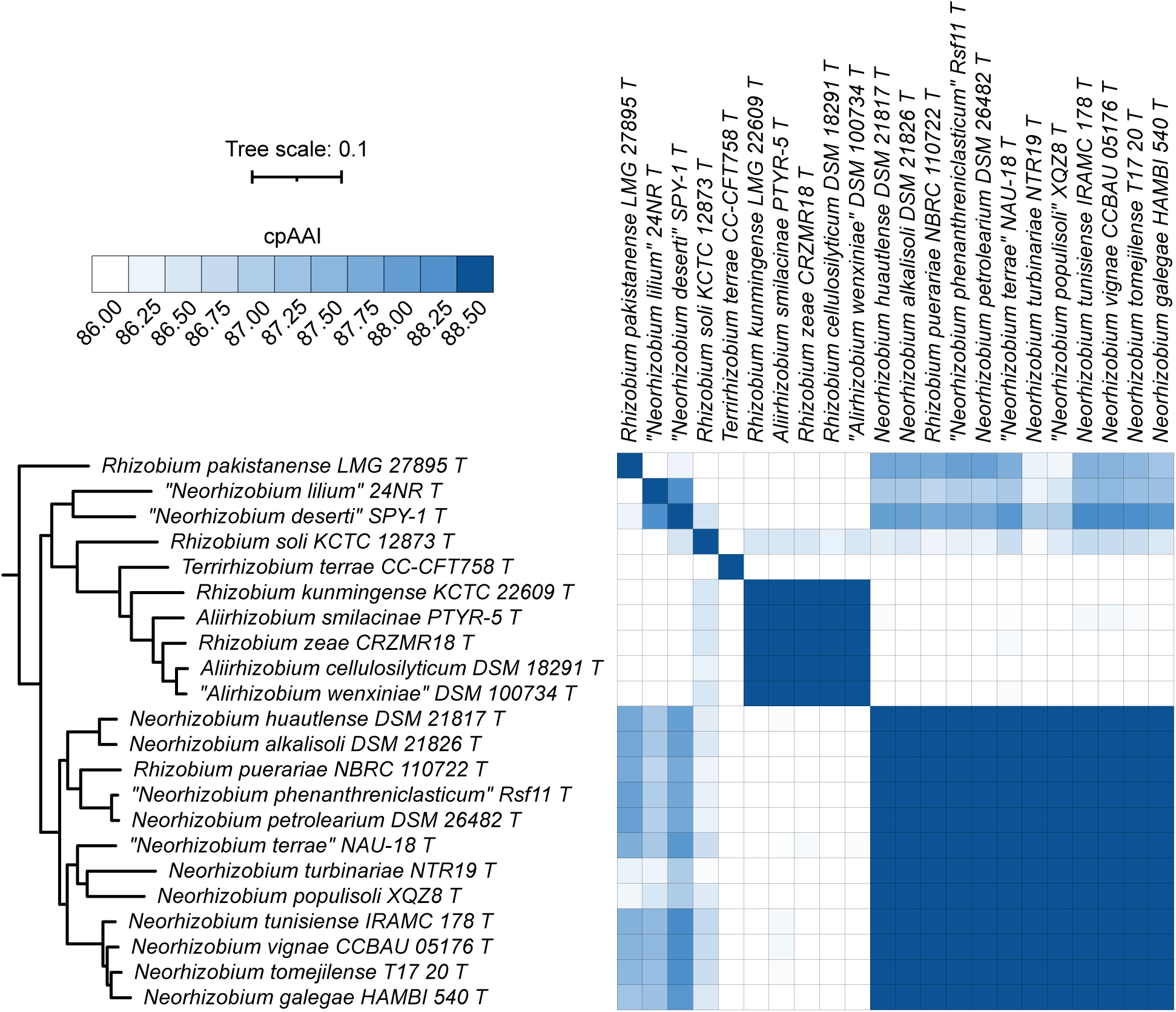
Phylogenomic analysis of the genera *Neorhizobium*, *Aliirhizobium*, and *Terrirhizobium*. (**Left**) A maximum likelihood phylogeny of the genera *Neorhizobium*, *Aliirhizobium*, and *Terrirhizobium*, together with related species; this phylogeny was extracted from the full *Rhizobiaceae* phylogeny of Figure S2. All support values calculated by SH-aLRT and ultra-fast jackknife were 100 and are therefore not shown. The scale bar represents the average number of amino acid substitutions per site. (**Right**) A matrix showing the core proteome average amino acid identity (cpAAI) values between each pair of strains calculated using a core set of 170 proteins. Values less than 86% are in white while all values greater than 88.5% are in the same shade of blue. An interactive version of this phylogeny is available at itol.embl.de/shared/1ps8ayxRcrNDU.

Based on the wide range of cpAAI values between the species type strains of this clade, we do not consider it appropriate to unify the genera *Neorhizobium*, *Aliirhizobium*, and *Terrirhizobium*. Instead, considering the cpAAI values, the topology of the maximum likelihood phylogeny, and a need to ensure each genus is monophyletic, we argue that *Rhizobium puerariae* should be reclassified as *Neorhizobium puerariae* comb. nov., *Rhizobium zeae* be reclassified as *Aliirhizobium zeae* comb. nov., *Rhizobium kunmingense* be reclassified as *Aliirhizobium kunmingense* comb. nov., and *R. soli* be reclassified as *Velazquezia soli* gen. nov. comb. nov.. Formal descriptions are provided below for those species (*N. puerariae* comb. nov., *A. zeae* comb. nov., *V. soli* gen. nov. comb. nov.) whose type strains currently meet the availability criteria specified by the ICNP. In addition, “*N. lilium*” and “*N. deserti*” should also be transferred to their own novel genus, while *R. pakistanense* should be transferred to a separate novel genus; however, these transfers cannot be formally proposed as none of these three type strains currently meet the availability criteria specified by the ICNP.

### Other taxonomic implications

Naranjo et al. [33] recently described the species *Agrobacterium divergens*. In our dataset, *A. divergens* R-31762^T^ is the deepest branching type strain of the genus *Agrobacterium*, and the cpAAI values with the other *Agrobacterium* type strains range from 84.5% to 86.6% (**Figure S2**, **Dataset S4**). Thus, it could reasonably be argued that *A. divergens* belongs to a novel genus. This argument is further supported by the fact that *A. divergens* strains lack the linear chromid that is a defining feature of all other members of the genus *Agrobacterium* [34, 35]. While we currently refrain from making this proposal, we argue that this reclassification would be justified if additional species that cluster with *A. divergens* are identified and if these species also lack linear chromids.

Lastly, our data are consistent with the transfer of “*Rhizobium halophilum*” to the genus *Pseudorhizobium*, “*Rhizobium quercicola*” to the genus *Ferranicluibacter*, and “*Rhizobium alarense*” to a novel genus (**Figure S2**, **Dataset S4**). However, as none of the three proposed type strains currently meet the availability criteria specified by the ICNP, we are unable to formally make these proposals.

Description of *Allohoeflea* gen. nov.

*Allohoeflea* (Al.lo.hoe’fle.a.; Gr. masc. pron. *allos*, other; N.L. fem. n. *Hoeflea*, a bacterial genus name; N.L. fem. n. *Allohoeflea*, the other Hoeflea).

The description is as given for *Allohoeflea poritis* comb. nov., which is the type species. The genus *Allohoeflea* has been separated from other *Rhizobiaceae* genera based on a core-proteome phylogeny and cpAAI values.

Description of *Arminia* gen. nov.

*Arminia* (Ar.mi’ni.a.; N.L. fem. n. *Arminia*, named to honour Professor Armin Braun, who established the concept of a tumor-inducing principle in crown-gall research).

Cells are Gram-negative, non-spore-forming rods, motile, and aerobic. The major fatty acids include C_19:0_ cyclo ω8c, C_18:1_ ω7c, C_18:1_ ω7c 11Me, C_16: 0_, C_14:0_ 3OH. The G+C content as calculated from genome sequences is around 59.3-60.0mol%. The type species is *Arminia tumorigenes* comb. nov..

Description of *Fluviimicrobium* gen. nov.

*Fluviimicrobium* (Flu.vi.i.mi.cro’bi.um.; L. masc. n. *fluvius*, a river; N.L. neut. n. *microbium*, a microbe; N.L. neut. N. *Fluviimicrobium*, a microbe from a river).

The description is as given for *Fluviimicrobium alvei* comb. nov., which is the type species. The genus *Fluviimicrobium* has been separated from other *Rhizobiaceae* genera based on a core-proteome phylogeny and cpAAI values.

Description of *Gillisella* gen. nov.

*Gillisella* (Gil.li.sel’la.; N.L. fem. n. *Gillisella*, named to honour Professor Monique Gillis, who made important contributions to the taxonomy of rhizobia).

Cells are Gram-negative, non-spore-forming rods, motile, and aerobic. The major ubiquinone is Q-10. The major fatty acids include C_19:0_ cyclo ω8c, C_18 :1_ ω7c, C_18 : 1_ 2-OH, C_16 : 0_, and C_16:0_ 3OH. The G+C content as calculated from genome sequences is around 55.0-59.3mol%. The type species is *Gillisella vitis* comb. nov..

Description of *Limnomicrobium* gen. nov.

*Limnomicrobium* (Lim.no.mi.cro’bi.um.; Gr. Fem. n. *limnê*, pool of standing water, lake; N.L. neut. N. *microbium*, a microbe; N.L. neut. n. *Limnomicrobium*, a microbe from a lake).

The description is as given for *Limnomicrobium aquaticum* comb. nov., which is the type species. The genus *Limnomicrobium* has been separated from other *Rhizobiaceae* genera based on a core-proteome phylogeny and cpAAI values.

Description of *Martinezia* gen. nov.

*Martinezia* (Mar.ti.ne′zi.a.; N.L. fem. n. *Martinezia*, named to honour Professor Esperanza Martínez-Romero, who has made important contributions to the taxonomy of rhizobia).

Cells are Gram-negative, non-spore-forming rods, motile, and aerobic. The major fatty acids include C_19:0_ cyclo ω8c, C_18 :1_ ω7c, C_18:0_, C_17;1_ω8c, C_16 : 0_, C_16:0_ 3OH, and iso-C_15:0_ 3-OH. The G+C content as calculated from genome sequences is around 59.1-59.9mol%. The type species is *Martinezia tropici* comb. nov..

Description of *Neohoeflea* gen. nov.

*Neohoeflea* (Ne.o.hoe’fle.a.; Gr. masc. pron. *neos*, new; N.L. fem. n. *Hoeflea*, a bacterial genus name; N.L. fem. n. *Neohoeflea*, the new Hoeflea).

The description is as given for *Neohoeflea coraliihabitans* comb. nov., which is the type species. The genus *Neohoeflea* has been separated from other *Rhizobiaceae* genera based on a core-proteome phylogeny and cpAAI values.

Description of *Parahoeflea* gen. nov.

*Parahoeflea* (Pa.ra.hoe’fle.a.; Gr. prep. *para*, beside, alongside of; N.L. fem. n. *Hoeflea*, a bacterial genus name; N.L. fem. n. *Parahoeflea*, a genus adjacent to *Hoeflea*).

Cells are Gram-negative, non-spore-forming rods, motile, and aerobic. The major ubiquinone is Q-10. The major fatty acids include C_18 :1_ ω7c, C_18 :1_ ω7c 11Me, C_18:0_, C_16:1_ ω7c, and C_16 : 0_. The G+C content as calculated from genome sequences is around 59.8-65.6mol%. The type species is *Parahoeflea phototrophica* comb. nov..

Description of *Velazquezia* gen. nov.

*Velazquezia* (Ve.laz.que′zi.a.; N.L. fem. n. *Velazquezia*, named to honour Professor Encarna Velázquez, who has made important contributions to the taxonomy of rhizobia).

The description is as given for *Velazquezia soli* comb. nov., which is the type species. The genus *Velazquezia* has been separated from other *Rhizobiaceae* genera based on a core-proteome phylogeny and cpAAI values.

Description of *Yannia* gen. nov.

*Yannia* (Yan’ni.a.; N.L. fem. n. *Yannia*, named to honour Professor Youssef Yanni who studied the interactions of *Rhizobium leguminosarum* with rice and clover).

The description is as given for *Yannia setariae* comb. nov., which is the type species. The genus *Yannia* has been separated from other *Rhizobiaceae* genera based on a core-proteome phylogeny and cpAAI values.

Description of *Affinirhizobium capsici* comb. nov.

*Affinirhizobium capsici* (cap’si.ci.; N.L. gen. neut. n. *capsici*, of Capsicum, referring to the isolation of the type strain from tumors of Capsicum annuum var. grossum L.).

Basonym: *Rhizobium capsici* Lin *et al*. 2015.

The description is as provided by Lin *et al*. 2015 [36]. *A. capsici* can be differentiated from other

*Affinirhizobium* species based on ANI calculations. The genomic G+C content of the type strain is 56.0mol%, and its approximate genome size is 4.53 Mbp.

The type strain is CC-SKC2^T^ (=JCM 19535^T^ =BCRC 80699^T^). The ENA accession number for the genome sequence is ERZ28549281.

Description of Affinirhizobium straminoryzae comb. nov.

*Affinirhizobium straminoryzae* (stra.mi.no.ry’zae.; L. neut. n. *stramen (gen. straminis)*, straw; L. fem. n. *oryza*, rice; N.L. gen. fem. n. *straminoryzae*, of rice straw). Basonym: *Rhizobium straminoryzae* Lin *et al*. 2014.

The description is as provided by Lin *et al*. 2014 [37]. *A. straminoryzae* can be differentiated from other *Affinirhizobium* species based on ANI calculations. The genomic G+C content of the type strain is 63.7mol%, and its approximate genome size is 5.26 Mbp.

The type strain is CC-LY845^T^ (=JCM 19536^T^ =BCRC 80698^T^). The ENA accession number for the genome sequence is ERZ28549277.

Description of *Aliirhizobium zeae* comb. nov.

*Aliirhizobium zeae* (ze’ae.; L. gen. fem. n. *zeae*, of *Zea mays*). Basonym: *Rhizobium zeae* Celador-Lera *et al*. 2017.

The description is as provided by Celador-Lera *et al*. 2017 [38]. *A. zeae* can be differentiated from other *Aliirhizobium* species based on ANI calculations. The genomic G+C content of the type strain is 59.4mol%, and its approximate genome size is 6.27 Mbp.

The type strain is CRZM18R^T^ (=LMG 29735^T^ =CECT 9169^T^). The ENA accession number for the genome sequence is ERZ28549278.

Description of *Allohoeflea poritis* comb. nov.

*Allohoeflea poritis* (po.ri’tis.; N.L. gen. fem. n. *poritis*, of the coral genus *Porites*).

Basonym: *Hoeflea poritis* Zhang *et al*. 2023.

The description is as provided by Zhang et al. 2023 [39]. *A. poritis* can be differentiated from other *Rhizobiaceae* species based on ANI calculations. The genomic G+C content of the type strain is 60.3mol%, and its approximate genome size is 5.58 Mbp.

The type strain is E7-10^T^ (=JCM 35852^T^ =MCCC 1K08229^T^). The NCBI RefSeq assembly accession number for the genome sequence is GCF_027889715.1.

Description of *Arminia rhododendri* comb. nov.

*Arminia rhododendri* (rho.do.den’dri.; N.L. gen. neut. n. *rhododendri*, of *Rhododendron*, the plant genus from which the type strain was isolated).

Basonym: *Rhizobium rhododendri* Kuzmanović et al. 2023.

The description is as provided by Kuzmanović et al. 2023 [29]. *A. rhododendri* can be differentiated from other *Arminia* species based on ANI calculations. The genomic G+C content of the type strain is 60.0mol%, and its approximate genome size is 5.96 Mbp.

The type strain is rho-6.2^T^ (=CFBP 9067^T^ = DSM 110655^T^). The NCBI RefSeq assembly accession number for the genome sequence is GCF_007000325.2.

Description of *Arminia tubonensis* comb. nov.

*Arminia tubonensis* (tu.bo.nen’sis.; N.L. fem. adj. *tubonensis*, pertaining to the ancient name "Tubo" of Tibet, where the bacterium was isolated).

Basonym: *Rhizobium tubonense* Zhang et al. 2011.

The description is as provided by Zhang et al. 2011 [30]. *A. tubonensis* can be differentiated from other *Arminia* species based on ANI calculations. The genomic G+C content of the type strain is 59.3mol%, and its approximate genome size is 6.54 Mbp.

The type strain is CCBAU 85046^T^ (=DSM 25379^T^ =HAMBI 3066^T^ =LMG 25225^T^). The NCBI RefSeq assembly accession number for the genome sequence is GCF_003240585.1.

Description of *Arminia tumorigenes* comb. nov.

*Arminia tumorigenes* (tu.mo.ri’ge.nes.; L. masc. n. *tumor*, swelling, tumor; Gr. suff. *-genes*, producing; from Gr. ind. v. *gennaô*, to produce; N.L. neut. part. adj. *tumorigenes*, tumor-producing).

Basonym: *Rhizobium tumorigenes* Kuzmanović et al. 2018.

The description is as provided by Kuzmanović et al. 2018 [31]. *A. tumorigenes* can be differentiated from other *Arminia* species based on ANI calculations. The genomic G+C content of the type strain is 60.0mol%, and its approximate genome size is 5.98 Mbp.

The type strain is 1078^T^ (=CFBP 8567^T^ = DSM 104880^T^). The NCBI RefSeq assembly accession number for the genome sequence is GCF_003240565.2.

Description of *Fluviimicrobium alvei* comb. nov.

*Fluviimicrobium alvei* (al’ve.i.; L. gen. masc. n. *alvei*, of/from the bed of a river, referring to the location from where the type strain was isolated).

Basonym: *Rhizobium alvei* Sheu *et al*. 2015.

The description is as provided by Sheu *et al*. 2015 [40]. *F. alvei* can be differentiated from other *Rhizobiaceae* species based on ANI calculations. The genomic G+C content of the type strain is 59.6mol%, and its approximate genome size is 5.20 Mbp.

The type strain is TNR-22^T^ (=BCRC 80408^T^ =DSM 100976^T^ =KCTC 23919^T^ =LMG 26895^T^). The NCBI RefSeq assembly accession number for the genome sequence is GCF_030551875.1.

Description of *Gillisella ampelina* comb. nov.

*Gillisella ampelina* (am.pe’li.na.; Gr. fem. n. *ampelos*, grapevine; Gr. masc./fem. adj. *ampelinos*; L. fem. adj. *ampelina*, of the vine).

Basonym: *Allorhizobium ampelinum* Kuzmanović et al. 2022.

The description is as provided by Kuzmanović et al. 2022 [41]. *G. ampelina* can be differentiated from other *Gillisella* species based on ANI calculations. The genomic G+C content of the type strain is 57.5mol%, and its approximate genome size is 6.32 Mbp.

The type strain is S4^T^ (=ATCC BAA-846^T^ =DSM 112012^T^). The NCBI RefSeq assembly accession number for the genome sequence is GCF_000016285.1.

Description of Gillisella taibaishanensis comb. nov.

*Gillisella taibaishanensis* (tai.bai.sha.nen’sis.; N.L. fem. adj. *taibaishanensis*, of or belonging to the Taibaishan Mountains in the Shaanxi province of China, where the bacterium was isolated).

Basonym: *Allorhizobium taibaishanense* (Yao *et al*. 2012) Hördt *et al*. 2020.

The description is as provided by Hördt et al. 2020 [5]. *G. taibaishanensis* can be differentiated from other *Gillisella* species based on ANI calculations. The genomic G+C content of the type strain is 59.3mol%, and its approximate genome size is 5.42 Mbp.

The type strain is 14971^T^ (=ACCC 14971^T^ =CCNWSX 483^T^ =DSM 100021^T^ =HAMBI 3214^T^). The NCBI RefSeq assembly accession number for the genome sequence is GCF_001938985.1.

Description of *Gillisella terrae* comb. nov.

*Gillisella terrae* (ter’rae.; L. gen. fem. n. *terrae*, of/from soil).

Basonym: *Allorhizobium terrae* Lin et al. 2020.

The description is as provided by Lin et al. 2020 [42]. *G. terrae* can be differentiated from other *Gillisella* species based on ANI calculations. The genomic G+C content of the type strain is 55.0mol%, and its approximate genome size is 4.50 Mbp.

The type strain is CC-HIH110^T^ (=BCRC 80932^T^ =JCM 31228^T^). The NCBI RefSeq assembly accession number for the genome sequence is GCF_004801395.1.

Description of *Gillisella vitis* comb. nov.

*Gillisella vitis* (vi’tis.; L. fem. n. *Vitis*, wine plant; L. gen. fem. n. *vitis*, of the wine plant).

Basonym: *Allorhizobium vitis* (Ophel and Kerr 1990) Mousavi *et al*. 2015.

The description is as provided by Mousavi et al. 2015 [43]. *G. vitis* can be differentiated from other *Gillisella* species based on ANI calculations. The genomic G+C content of the type strain is 57.6mol%, and its approximate genome size is 5.75 Mbp.

The type strain is K309^T^ (=ATCC 49767^T^ =CECT 4799^T^ =CIP 105853^T^ =HAMBI 1817^T^ =ICMP 10752^T^ =IFO 15140^T^ =JCM 21033^T^ =LMG 8750^T^ =NBRC 15140^T^ =NCPPB 3554^T^). The NCBI RefSeq assembly accession number for the genome sequence is GCF_001541345.2.

Description of *Limnomicrobium aquaticum* comb. nov.

*Limnomicrobium aquaticum* (a.qua’ti.cum.; L. neut. adj. *aquaticum*, living in water, aquatic, referring to the isolation source of the type strain).

Basonym: *Rhizobium aquaticum* Máthé *et al*. 2018.

The description is as provided by Máthé *et al*. 2018 [44]. *L. aquaticum* can be differentiated from other *Rhizobiaceae* species based on ANI calculations. The genomic G+C content of the type strain is 61.0mol%, and its approximate genome size is 4.91 Mbp.

The type strain is SA-276^T^ (=DSM 29780^T^ = JCM 31760^T^). The NCBI RefSeq assembly accession number for the genome sequence is GCF_040545545.1.

Description of *Martinezia calliandrae* comb. nov.

*Martinezia calliandrae* (cal.li.an’drae.; N.L. gen. fem. n. *calliandrae*, of *Calliandra*, the genus of the medicinal plant *C. grandiflora* from which bacteria were isolated).

Basonym: *Rhizobium calliandrae* Rincón-Rosales et al. 2013.

The description is as provided by Rincón-Rosales et al. 2013 [45]. *M. calliandrae* can be differentiated from other *Martinezia* species based on ANI calculations. The genomic G+C content of the type strain is 59.1mol%, and its approximate genome size is 7.55 Mbp.

The type strain is LBP2-1^T^ (=ATCC BAA-2435^T^ =CCGE 524^T^ =CIP 110456^T^). The NCBI RefSeq assembly accession number for the genome sequence is GCF_030272695.1.

Description of *Martinezia dioscoreae* comb. nov.

*Martinezia dioscoreae* (di.os.co.re’ae.; N.L. gen. fem. n. *dioscoreae*, of the plant genus *Dioscorea* species).

Basonym: *Rhizobium dioscoreae* Ouyabe et al. 2020.

The description is as provided by Ouyabe et al. 2020 [46]. *M. dioscoreae* can be differentiated from other *Martinezia* species based on ANI calculations. The genomic G+C content of the type strain is 59.7mol%, and its approximate genome size is 6.73 Mbp.

The type strain is S-93^T^ (=DSM 110498^T^ =NBRC 114257^T^ =NRIC 988^T^). The NCBI RefSeq assembly accession number for the genome sequence is GCF_009176305.1.

Description of *Martinezia freirei* comb. nov.

*Martinezia freirei* (frei’re.i.; N.L. gen. masc. n. *freirei*, of Freire, named after Professor João Ruy Jardim Freire, a distinguished Brazilian rhizobiologist).

Basonym: *Rhizobium freirei* Dall’Agnol et al. 2013.

The description is as provided by Dall’Agnol et al. 2013 [47]. *M. freirei* can be differentiated from other *Martinezia* species based on ANI calculations. The genomic G+C content of the type strain is 59.7mol%, and its approximate genome size is 6.73 Mbp.

The type strain is PRF 81^T^ (=CNPSo 122^T^ =IPR-Pv81^T^ =LMG 27576^T^ =SEMIA 4080^T^ =WDCM 440^T^).

The NCBI RefSeq assembly accession number for the genome sequence is GCF_000359745.1.

Description of *Martinezia hainanensis* comb. nov.

*Martinezia hainanensis* (hai.na.nen’sis.; N.L. fem. adj. *hainanensis*, pertaining to Hainan Province in China).

Basonym: *Rhizobium hainanense* Chen et al. 1997.

The description is as provided by Chen et al. 1997 [48]. *M. hainanensis* can be differentiated from other *Martinezia* species based on ANI calculations. The genomic G+C content of the type strain is 59.7mol%, and its approximate genome size is 7.25 Mbp.

The type strain is I66^T^ (=CCBAU 57015^T^ =CIP 105503^T^ =DSM 11917^T^). The NCBI RefSeq assembly accession number for the genome sequence is GCF_900094555.1.

Description of *Martinezia jaguaris* comb. nov.

*Martinezia jaguaris* (ja.gu.a.ris.; N.L. gen. masc. n. *jaguaris*, jaguar). Basonym: *Rhizobium jaguaris* Rincón-Rosales et al. 2013.

The description is as provided by Rincón-Rosales et al. 2013 [45]. *M. jaguaris* can be differentiated from other *Martinezia* species based on ANI calculations. The genomic G+C content of the type strain is 59.4mol%, and its approximate genome size is 8.03 Mbp.

The type strain is SJP1-2^T^ (=ATCC BAA-2445^T^ =CCGE 525^T^ =CIP 110453^T^). The NCBI RefSeq assembly accession number for the genome sequence is GCF_003627755.1.

Description of *Martinezia leucaenae* comb. nov.

*Martinezia leucaenae* (leu.cae’nae.; N.L. gen. fem. n. *leucaenae*, of Leucaena, referring to the isolation source of many strains of this species, root nodules of *Leucaena*).

Basonym: *Rhizobium leucaenae* Ribeiro et al. 2012.

The description is as provided by Ribeiro et al. 2012 [49]. *M. leucaenae* can be differentiated from other *Martinezia* species based on ANI calculations. The genomic G+C content of the type strain is 59.4mol%, and its approximate genome size is 6.68 Mbp.

The type strain is CFN 299^T^ (=CECT 4844^T^ =CENA 183^T^ =CNPSo 141^T^ =IAM 14230^T^ =JCM 21088^T^ =LMG 9517^T^ =SEMIA 4083^T^ =UMR 1026^T^ =USDA 9039^T^). The NCBI RefSeq assembly accession number for the genome sequence is GCF_000426285.1.

Description of *Martinezia lusitana* comb. nov.

*Martinezia lusitana* (lu.si.ta’na.; N.L. fem. adj. *lusitana*, of Lusitania, the Roman name of Portugal, where the strains reported in this study were isolated).

Basonym: *Rhizobium lusitanum* Valverde et al. 2006.

The description is as provided by Valverde et al. 2006 [50]. *M. lusitana* can be differentiated from other *Martinezia* species based on ANI calculations. The genomic G+C content of the type strain is 59.7mol%, and its approximate genome size is 7.92 Mbp.

The type strain is P1-7^T^ (=CECT 7016^T^ =LMG 22705^T^). The NCBI RefSeq assembly accession number for the genome sequence is GCF_900094565.1.

Description of *Martinezia multihospitum* comb. nov.

*Martinezia multihospitum* (mul.ti.hos’pi.tum.; L. masc. adj. *multus*, many, numerous; L. masc. n. *hospes (gen. pl. hospitum)*, he who entertains a stranger, a host; N.L. gen. masc. pl. multihospitum, of numerous hosts, referring to the isolation of the bacterium from various legume species).

Basonym: *Rhizobium multihospitium* Han et al. 2008.

The description is as provided by Han et al. 2008 [51]. *M. multihospitum* can be differentiated from other *Martinezia* species based on ANI calculations. The genomic G+C content of the type strain is 59.8mol%, and its approximate genome size is 7.32 Mbp.

The type strain is CCBAU 83401^T^ (=DSM 21814^T^ =HAMBI 2975^T^ =LMG 23946^T^). The NCBI RefSeq assembly accession number for the genome sequence is GCF_900094585.1.

Description of *Martinezia miluonensis* comb. nov.

*Martinezia miluonensis* (mi.lu.o.nen’sis.; N.L. fem. adj. *miluonensis*, pertaining to the Miluo River, a famous river located in Hunan Province, where the bacterium was isolated).

Basonym: *Rhizobium miluonense* Gu et al. 2008.

The description is as provided by Gu et al. 2008 [52]. *M. miluonensis* can be differentiated from other *Martinezia* species based on ANI calculations. The genomic G+C content of the type strain is 59.7mol%, and its approximate genome size is 7.92 Mbp.

The type strain is CCBAU 41251^T^ (=DSM 21815^T^ =HAMBI 2971^T^ =LMG 24208^T^). The NCBI RefSeq assembly accession number for the genome sequence is GCF_900094545.1.

Description of *Martinezia paranaensis* comb. nov.

*Martinezia paranaensis* (pa.ra.na.en’sis.; N.L. fem. adj. *paranaensis*, of or belonging to Paraná. Named after Paraná State, where the type strain was isolated).

Basonym: *Rhizobium paranaense* Dall’Agnol et al. 2014.

The description is as provided by Dall’Agnol et al. 2014 [53]. *M. paranaensis* can be differentiated from other *Martinezia* species based on ANI calculations. The genomic G+C content of the type strain is 59.6mol%, and its approximate genome size is 6.86 Mbp.

The type strain is PRF 35^T^ (=CNPSo 120^T^ =IPR-Pv1249^T^ =LMG 27577^T^). The ENA accession number for the genome sequence is ERZ28549275.

Description of *Martinezia rhizogenes* comb. nov.

*Martinezia rhizogenes* (rhi.zo’ge.nes.; Gr. fem. n. *rhiza*, root; Gr. suff. *-genes*, producing; from Gr. ind. v. *gennaô*, to produce; N.L. fem. part. adj. *rhizogenes*, root-producing).

Basonym: *Rhizobium rhizogenes* (Riker *et al*. 1930) Young *et al*. 2001.

The description is as provided by Young et al. 2001 [54]. *M. rhizogenes* can be differentiated from other *Martinezia* species based on ANI calculations. The genomic G+C content of the type strain is 59.9mol%, and its approximate genome size is 7.05 Mbp.

The type strain is ATCC 11325^T^ (=CFBP 5520^T^ =CIP 104328^T^ =DSM 30148^T^ =HAMBI 1816^T^ =ICMP 5794^T^ =IFO 13257^T^ =JCM 20919^T^ =LMG 150^T^ =NBRC 13257^T^ =NCPPB 2991^T^). The NCBI RefSeq assembly accession number for the genome sequence is GCF_007002985.1.

Description of *Martinezia tropici* comb. nov.

*Martinezia tropici* (tro’pi.ci.; N.L. gen. masc. n. *tropici*, of the tropic (of Cancer); from L. masc. adj. *tropicus*, tropical).

Basonym: *Rhizobium tropici* Martínez-Romero et al. 1991.

The description is as provided by Martínez-Romero et al. 1991 [55]. *M. tropici* can be differentiated from other *Martinezia* species based on ANI calculations. The genomic G+C content of the type strain is 59.5mol%, and its approximate genome size is 6.69 Mbp.

The type strain is CIAT 899^T^ (=ATCC 49672^T^ =DSM 11418^T^ =HAMBI 1163^T^ =IFO 15247^T^ =JCM 21072^T^ =LMG 9503^T^ =NBRC 15247^T^). The NCBI RefSeq assembly accession number for the genome sequence is GCF_000330885.1.

Description of Neohoeflea coraliihabitans comb. nov.

*Neohoeflea coraliihabitans* (co.ra.li.i.ha’bi.tans.; N.L. neut. n. *coralium*, coral; L. pres. Part. *habitans*, inhabiting; N.L. fem. part. adj. *Coraliihabitans*, inhabiting the corals).

Basonym: *Pseudohoeflea coraliihabitans* Yu *et al*. 2024.

The description is as provided by Yu *et al*. 2024 [56]. *N. coraliihabitans* can be differentiated from other *Rhizobiaceae* species based on ANI calculations. The genomic G+C content of the type strain is 62.7mol%, and its approximate genome size is 3.93 Mbp.

The type strain is DPN4N28-3^T^ (=KCTC 82803^T^ =MCCC 1K05639^T^). The NCBI RefSeq assembly accession number for the genome sequence is GCF_019334345.1.

Description of *Neorhizobium puerariae* comb. nov.

*Neorhizobium puerariae* (pu.e.ra’ri.ae.; N.L. gen. fem. n. *puerariae*, of *Pueraria*, the genus of the plant *Pueraria candollei*, from which the type strain was isolated).

Basonym: *Rhizobium puerariae* Boonsnongcheep *et al*. 2016.

The description is as provided by Boonsnongcheep *et al*. 2016 [57]. *N. puerariae* can be differentiated from other *Neorhizobium* species based on ANI calculations. The genomic G+C content of the type strain is 62.7mol%, and its approximate genome size is 6.63 Mbp.

The type strain is PC004^T^ (=BCC 73740^T^ =NBRC 110722^T^). The ENA accession number for the genome sequence is ERZ28549276.

Description of *Paenirhizobium cremeum* comb. nov.

*Paenirhizobium cremeum* (cre.me’um.; N.L. neut. adj. *cremeum*, cream-white, referring to the colour of the colonies).

Basonym: *Rhizobium cremeum* Yang *et al*. 2022.

The description is as provided by Yang *et al*. 2022 [58]. *P. cremeum* can be differentiated from other *Paenirhizobium* species based on ANI calculations. The genomic G+C content of the type strain is 61.6mol%, and its approximate genome size is 5.33 Mbp.

The type strain is W15^T^ (=CGMCC 1.18731^T^ =KACC 22344^T^). The NCBI RefSeq assembly accession number for the genome sequence is GCF_022884065.1.

Description of *Parahoeflea alexandrii* comb. nov.

*Parahoeflea alexandrii* (a.le.xan’dri.i.; N.L. gen. neut. n. *alexandrii*, of *Alexandrium*, the genus name of the dinoflagellate *Alexandrium minutum*, the source of isolation of the type strain).

Basonym: *Hoeflea alexandrii* Palacios *et al*. 2006.

The description is as provided by Palacios *et al*. 2006 [59]. *P. alexandrii* can be differentiated from other *Parahoeflea* species based on ANI calculations. The genomic G+C content of the type strain is 61.6mol%, and its approximate genome size is 4.92 Mbp.

The type strain is AM1V30^T^ (=CECT 5682^T^ =DSM 16655^T^). The NCBI RefSeq assembly accession number for the genome sequence is GCF_024105735.1.

Description of *Parahoeflea halophila* comb. nov.

*Parahoeflea halophila* (ha.lo’phi.la.; Gr. masc. n. *hals (gen. halos)*, salt; N.L. fem. adj. suff. -*phila*, friend, loving; from Gr. fem. adj. *philê*, loving; N.L. fem. adj. *halophila*, salt-loving).

Basonym: *Hoeflea halophila* Jung *et al*. 2013.

The description is as provided by Jung *et al*. 2013 [60]. *P. halophila* can be differentiated from other *Parahoeflea* species based on ANI calculations. The genomic G+C content of the type strain is 61.1mol%, and its approximate genome size is 4.19 Mbp.

The type strain is JG120-1^T^ (=JCM 16715^T^ =KCTC 23107^T^). The NCBI RefSeq assembly accession number for the genome sequence is GCF_900220985.1.

Description of *Parahoeflea olei* comb. nov.

*Parahoeflea olei* (o’le.i.; L. gen. neut. n. *olei*, of oil). Basonym: *Hoeflea olei* Rahul *et al*. 2015.

The description is as provided by Rahul *et al*. 2015 [61]. *P. olei* can be differentiated from other *Parahoeflea* species based on ANI calculations. The genomic G+C content of the type strain is 65.6mol%, and its approximate genome size is 4.72 Mbp.

The type strain is JC230^T^ (=KCTC 42071^T^ =LMG 28200^T^). The NCBI RefSeq assembly accession number for the genome sequence is GCF_001703635.1.

Description of Parahoeflea phototrophica comb. nov.

*Parahoeflea phototrophica* (pho.to.tro’phi.ca.; Gr. neut. n. *phôs (gen. phôtos)*, light; Gr. masc. adj. *trophikos*, nursing, tending or feeding; N.L. fem. adj. *phototrophica*, referring to the likely ability to use light for energy generation).

Basonym: *Hoeflea phototrophica* Biebl *et al*. 2006.

The description is as provided by Biebl *et al*. 2006 [62]. *P. phototrophica* can be differentiated from other *Parahoeflea* species based on ANI calculations. The genomic G+C content of the type strain is 59.8mol%, and its approximate genome size is 4.47 Mbp.

The type strain is DFL-43^T^ (=DSM 17068^T^ =NCIMB 14078^T^). The NCBI RefSeq assembly accession number for the genome sequence is GCF_000154705.2.

Description of *Sinorhizobium aridi* comb. nov.

*Sinorhizobium aridi* (a’ri.di.; L. neut. n. *aridum* (gen. sing. *aridi*), a dry place, referring to the areas where bacteria were isolated from and which are subjected to droughts).

Basonym: *Ensifer aridi* Rocha *et al*. 2020.

The description is as provided by Rocha *et al*. 2020 [63]. *S. aridi* can be differentiated from other *Sinorhizobium* species based on ANI calculations. The genomic G+C content of the type strain is 61.8mol%, and its approximate genome size is 6.60 Mbp.

The type strain is LMR001^T^ (=HAMBI 3707^T^ =LMG 31426^T^). The NCBI RefSeq assembly accession number for the genome sequence is GCF_002078505.1.

Description of *Velazquezia soli* comb. nov.

*Velazquezia soli* (so’li.; L. gen. neut. n. *soli*, of soil). Basonym: *Rhizobium soli* Yoon *et al*. 2010.

The description is as provided by Yoon *et al*. 2010 [64]. *V. soli* can be differentiated from other *Rhizobiaceae* species based on ANI calculations. The genomic G+C content of the type strain is 60.0mol%, and its approximate genome size is 5.19 Mbp.

The type strain is DS-42^T^ (=JCM 14591^T^ =KCTC 12873^T^). The ENA accession number for the genome sequence is ERZ28549283.

Description of *Yannia setariae* comb. nov.

*Yannia setariae* (se.ta’ri.ae.; N.L. gen. fem. n. *setariae*, of *Setaria*, referring to the plant from which the type strain was isolated).

Basonym: *Rhizobium setariae* Kang and Seo 2022.

The description is as provided by Kang and Seo 2022 [65]. *Y. setariae* can be differentiated from other *Rhizobiaceae* species based on ANI calculations. The genomic G+C content of the type strain is 59.3mol%, and its approximate genome size is 5.61 Mbp.

The type strain is KVB221^T^ (=KACC 21713^T^ =NBRC 114644^T^). The NCBI RefSeq assembly accession number for the genome sequence is GCF_016722925.1.

## ACKNOWLEDGEMENTS

This work was supported by the Natural Sciences and Engineering Research Council of Canada (NSERC) through an Alliance International Catalyst grant (ALLRP-597321-24) to GCD, EM, and JPWY, as well as Grant PID2022-138373NA-I00 to EM and GCD, which is funded by MCIN/AEI/10.13039/501100011033 and by “ERDF A way of making Europe”. This research was also supported by the “Bio-inoculants for the promotion of nutrient use efficiency and crop resiliency in Canadian agriculture” project funded by the Government of Canada through Genome Canada and Genome Prairie (CSAFS-ICT 19308), and the Government of Ontario through an Ontario Research Fund – Interdisciplinary Challenge Teams (ORF-ICT) – Climate Action Genomics Initiative grant (File ICT 19308). In addition, EM acknowledges support from the ‘Escalera de Excelencia’ CLU-2025-2-04 program of the Regional Government of Castilla y León, co-funded by the Castilla y León 2021–2027 Operational Program (FEDER), Spain.

