## Supplementary_Materials for "Updated taxonomy of the family *Rhizobiaceae* with proposals for 10 novel genera and 35 novel combinations"

**This PDF file includes:**

Figure S1 (page 2)
Legend for Figure S2 (page 3)
Legends for Datasets S1 to S4 (page 4)

**Other supplementary materials for this manuscript include the following:**

Figure S2
Datasets S1 to S4

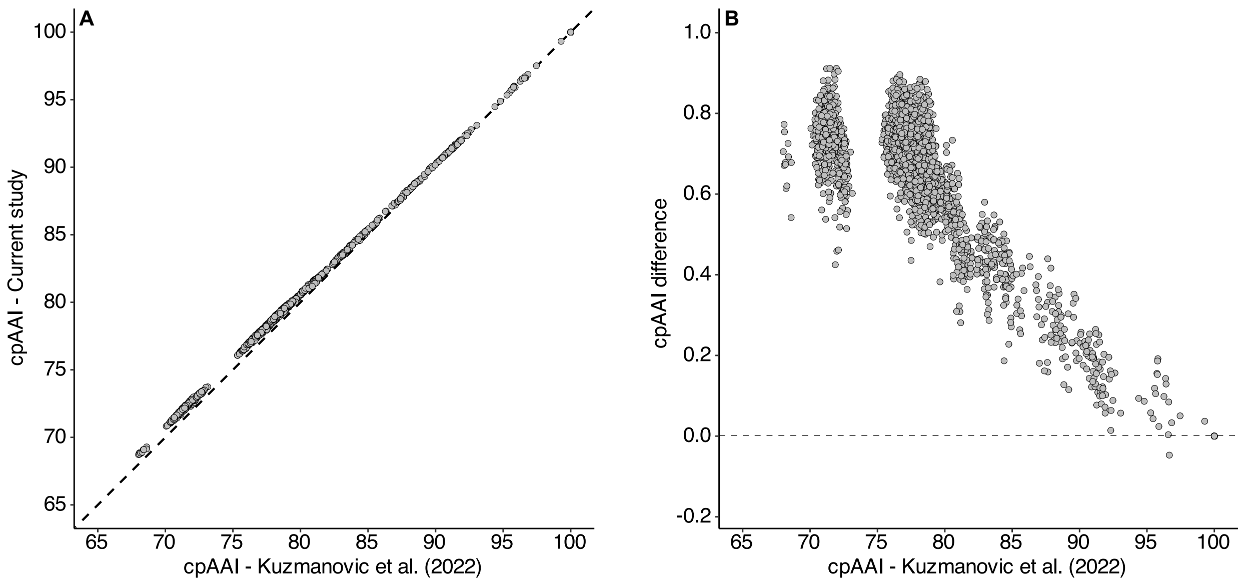


**Figure S1. Comparison of the pairwise cpAAI values calculated by Kuzmanović et al. (2022) and this study.** The pairwise core-proteome average amino acid identity (cpAAI) values calculated in this study and by Kuzmanović et al. (2022) were compared. (**A**) Each dot represents the cpAAI values for a species type strain comparison shared between studies, while the dashed horizontal line show the 1:1 relationship. (**B**) The difference in cpAAI values (calculated as cpAAI [current study] minus cpAAI [Kuzmanović et al. (2022)]) is plotted against to the cpAAI values calculated from Kuzmanović et al. (2022). The dashed line shows the 0 value.

**Kuzmanović N, Fagorzi C, Mengoni A, Lassalle F, diCenzo GC**. Taxonomy of Rhizobiaceae revisited: proposal of a new framework for genus delimitation. *Int J Syst Evol Microbiol* 2022;72:005243.

**Figure S2. Phylogenomic analysis of the family *Rhizobiaceae*.** (Figure provided as a separate file due to its size.) A maximum likelihood phylogeny of 242 type strains from the family *Rhizobiaceae*, rooted using three type strains from the genus *Mesorhizobium* (family *Bartonellaceae*). The numbers on the branches indicate support values by SH-aLRT (left) and ultra-fast jackknife (right); values are only shown at nodes where at least one value is below 100. The scale bar represents the average number of amino acid substitutions per site. The columns to the right of the phylogeny indicate the previous genus assignment (left column) and the proposed genus assignment (right column) of each species. In the rightmost column, boxes with a red outline indicate taxonomic changes that should be made but that cannot be made either because the original name is not yet validated or the type strain no longer meets the availability criteria specified by the ICNP. An interactive version of this phylogeny is available at itol.embl.de/shared/1ps8ayxRcrNDU.

### DATASET LEGENDS

**Dataset S1. Type strains sequenced as part of this study.** This dataset includes information for the 18 type strains sequenced as part of this study and includes both the current species name and, where relevant, the proposed species name for each type strain. In addition, ENA accessions are provided to facilitate access to the genome sequence and Oxford Nanopore Technologies read set for each type strain.

**Dataset S2. Type strains whose genomes were downloaded from NCBI.** This dataset includes information for the 222 type strains whose genomes were downloaded from NCBI, and includes both the current species name and, where relevant, the proposed species name for each type strain. In addition, the NCBI RefSeq or GenBank accessions to access the genomes of the type strains are provided.

**Dataset S3. Type strains whose genomes were downloaded from JGI.** This dataset includes information for the five type strains whose genomes were downloaded from JGI, and includes both the current species name and, where relevant, the proposed species name for each type strain. In addition, the JGI GOLD accessions to access the genomes of the type strains are provided.

**Dataset S4. Core-proteome average amino acid identity (cpAAI) matrix.** This dataset includes an all-against-all matrix showing the pairwise cpAAI values calculated between each pair of the 245 type strains analyzed in this study.
