## Supplementary figures and images for "Updated taxonomy of the family *Rhizobiaceae* with proposals for 10 novel genera and 35 novel combinations"

### Figure_S2

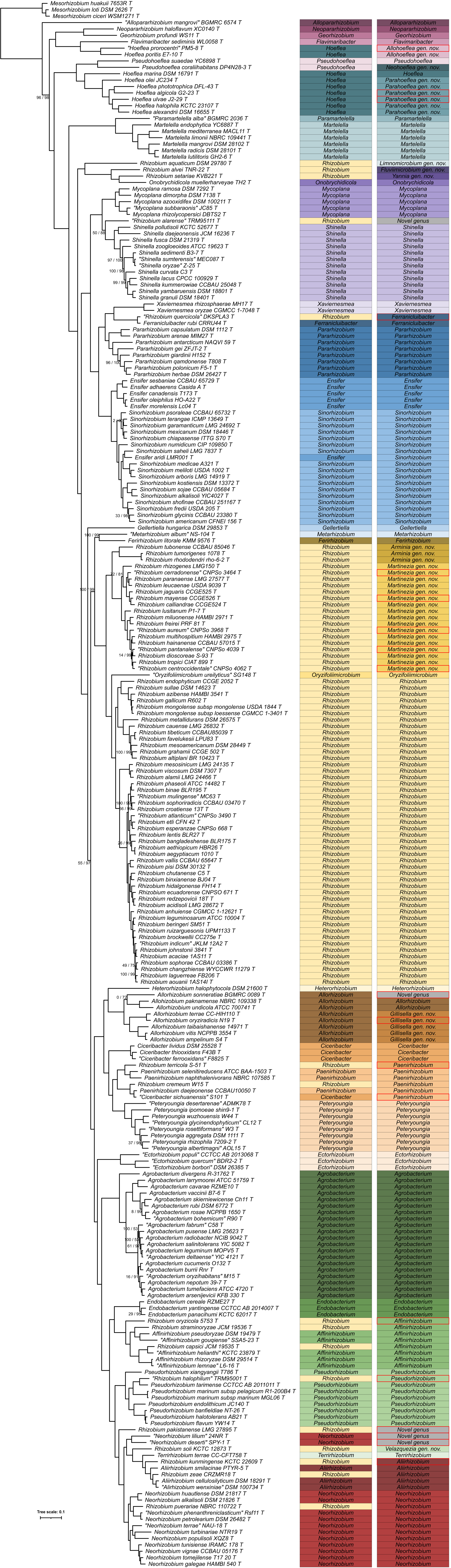
